# A recurring YYDRxG pattern in broadly neutralizing antibodies to a conserved site on SARS-CoV-2, variants of concern, and related viruses

**DOI:** 10.1101/2021.12.15.472864

**Authors:** Hejun Liu, Chengzi I. Kaku, Ge Song, Meng Yuan, Raiees Andrabi, Dennis R. Burton, Laura M. Walker, Ian A. Wilson

**Author notes:** Corresponding authors (I.A.W.), (L.M.W.). These authors contributed equally (Hejun Liu, Chengzi I. Kaku).

## Abstract

Studying the antibody response to SARS-CoV-2 informs on how the human immune system can respond to antigenic variants as well as other SARS-related viruses. Here, we structurally and functionally characterized a potent human antibody ADI-62113 that also neutralizes SARS-CoV- 2 variants of concern and cross-reacts with many other sarbecoviruses. A YYDRxG motif encoded by *IGHD3-22* in CDR H3 facilitates targeting to a highly conserved epitope on the SARS-CoV-2 receptor binding domain. A computational search for a YYDRxG pattern in publicly available sequences identified many antibodies with broad neutralization activity against SARS-CoV-2 variants and SARS-CoV. Thus, the YYDRxG motif represents a common convergent solution for the human humoral immune system to counteract sarbecoviruses. These findings also suggest an epitope targeting strategy to identify potent and broadly neutralizing antibodies that can aid in the design of pan-sarbecovirus vaccines and antibody therapeutics.

**Short Summary:** Decryption of a recurrent sequence feature in anti-SARS-CoV-2 antibodies identifies how potent pan-sarbecovirus antibodies target a conserved epitope on the receptor binding domain.

## INTRODUCTION

The current pandemic of coronavirus disease 2019 (COVID-19) has fomented devasting sociological and global economic consequences. Although several effective vaccines have been rapidly developed, SARS-CoV-2, the etiological cause of COVID-19, is still raging throughout the world. The efficacy of these vaccines has been affected by viral antigenic drift that has led to enhanced infectivity as well as escape neutralizing antibodies elicited by SARS-CoV-2 infection and vaccination. A majority of the potent neutralizing antibodies target the receptor binding domain (RBD) of the spike protein. However, these antibodies are often susceptible to mutations at the receptor binding site (RBS), which are frequently found in circulating SARS-CoV-2 variants (R. E. Chen et al., 2021; Hoffmann et al., 2021; P. Wang, M. S. Nair, et al., 2021; Yuan, Huang, et al., 2021; Yuan, Liu, et al., 2021). Notwithstanding, a subset of broadly neutralizing antibodies (bnAbs) can target more conserved surfaces on the spike protein of the virus and can neutralize circulating variants of concern (VOC), variants of interest (VOI), and often other SARS-related viruses in the sarbecovirus family (Jette et al., 2021; Li et al., 2021; Liu et al., 2020; H. Liu et al., 2021; Pinto et al., 2020; Rappazzo et al., 2021; Starr, Czudnochowski, et al., 2021; Tortorici et al., 2021; Wec et al., 2020). Identification and characterization of such cross-neutralizing antibodies are therefore urgently needed to fight the current COVID-19 pandemic, as well as prepare for future potential zoonotic spillover.

Despite their broad spectrum of neutralization activity against SARS-CoV-2, circulating and emerging variants, as well as other related coronaviruses with high pandemic risk, potent cross-neutralizing antibodies are rarely isolated compared to SARS-CoV-2-specific neutralizing antibodies. To date, only a limited number of cross-neutralizing antibodies that target regions in spikes that are highly conserved across sarbecoviruses, which include the CR3022 site and the N343 proteoglycan site in the RBD, as well as the S2 domain (Jette et al., 2021; Jiang et al., 2021; Li et al., 2021; Liu et al., 2020; H. Liu et al., 2021; Martinez et al., 2021; Piccoli et al., 2020; Pinto et al., 2020; Rappazzo et al., 2021; Sakharkar et al., 2021; Sauer et al., 2021; Starr, Czudnochowski, et al., 2021; Tortorici et al., 2021; Wec et al., 2020; Yuan, Liu, et al., 2021; Zhou et al., 2020; Zhou et al., 2021). We and others have reported structures of cross-neutralizing antibodies targeting the CR3022 site with potent neutralization against SARS-CoV-2, circulating and emerging variants, and other sarbecoviruses (Jette et al., 2021; Liu et al., 2020; Tortorici et al., 2021). However, the frequency that these cross-neutralizing antibodies are elicited by SARS-CoV-2 infection and vaccination, and why they are so rarely isolated, remain to be determined. Here, we structurally characterized such a cross-neutralizing antibody ADI-62113. We identified a recurrent YYDRxG motif in ADI-62113 encoded by *IGHD3-22* in heavy-chain complementarity-determining region 3 (CDR H3) from comparative analysis of the crystal structure of another cross-neutralizing antibody COVA1-16. A computational search of publicly available sequences with a YYDRxG sequence pattern identified more such antibodies that broadly neutralize SARS-CoV-2 VOC and SARS-CoV pseudoviruses, suggesting a general mechanism harnessed by the human humoral immune system to combat sarbecoviruses. Such information is critical for next-generation vaccine design and evaluation, as well as discovery of more effective therapeutic antibodies with increased breadth. Our study further suggests such cross-neutralizing bnAbs can be rapidly identified from their sequence alone.

## RESULTS

### Antibody ADI-62113 cross-reacts with a broad spectrum of sarbecoviruses

ADI-62113 is a potent cross-neutralizing antibody isolated from a COVID-19 patient (Sakharkar et al., 2021). Immunoglobulin heavy variable gene *IGHV1-3* and kappa variable gene *IGKV1-33* encode its heavy and light chain, respectively, and these germline genes have not been reported in other SARS-CoV-2 cross-neutralizing antibodies to date. To assess antibody breadth, we expressed various sarbecovirus RBDs on the surface of yeast to characterize their binding kinetics with antibody ADI-62113. ADI-62113 binds with high affinity to a broad spectrum of sarbecoviruses including ACE2-utilizing viruses in clade 1 and non-ACE2-utilizing viruses in clade 2 (Figure 1A). Thus, its binding properties are highly favorable as a potential pan-sarbecovirus prophylactic or therapeutic.

**Figure 1.**
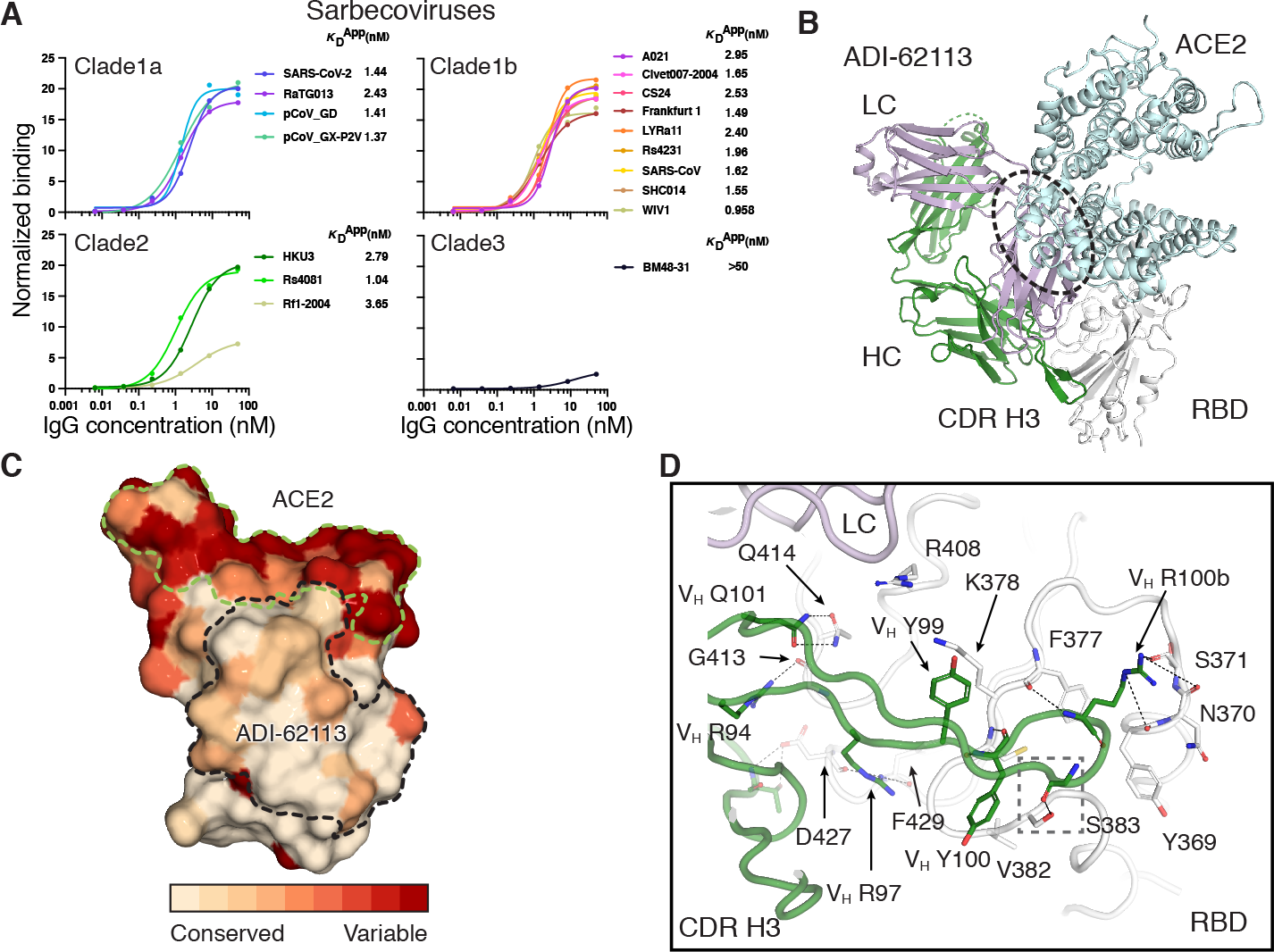
ADI-62113 binds a highly conserved site on SARS-CoV-2 RBD and cross-reacts with many sarbecoviruses. **A.** ADI-62113 shows a broad spectrum of cross-reactivity to sarbecoviruses. RBDs from viruses in clade 1a (SARS-CoV-2-like viruses), clade1b (SARS-CoV- like viruses), clade 2, and clade 3 were displayed on the surface of yeast for binding kinetics analysis. Clade 1a and 1b viruses use ACE2 as an entry receptor. Clade 2 and 3 viruses do not bind ACE2 but contain homologous sequences to SARS-CoV-2. **B.** Composite structure showing ADI-62113 competition with ACE2 binding for SARS-CoV-2 RBD. ADI-62113 Fab, ACE2, and SARS-CoV-2 RBD are shown in a ribbon representation. ACE2 is superimposed based on PDB ID: 6M0J. White, RBD; pale cyan, ACE2; dark green, heavy chain; lavender, light chain. ADI- 62113 binds to a highly conserved site on SARS-CoV-2 RBD, while the receptor binding site (RBS) is highly variable across sarbecoviruses. **C.** ADI-62113 binds a highly conserved surface. SARS-CoV-2 RBD is shown in surface representation and colored by conservation across sarbecovirus sequences. Dashed lines show the outline of the buried surface area (BSA) on the RBD by the human receptor ACE2 (green) and ADI-62113 (black). 81% of the ADI-62113 epitope surface is buried by the heavy chain and 19% by the light chain. CDR H3 contributes 69% of the total BSA. **D.** CDR H3 interacts with highly conserved residues in the RBD. The crystal structure of ADI- 62113 in complex with SARS-CoV-2 RBD is shown in tube representation. Residues involved in the interface between ADI-62113 and SARS-CoV-2 RBD are shown in sticks. Dashed lines represent hydrogen bonds or salt bridges. V_H_ R94, R97, R100b, G100d, and N101 hydrogen bond with the RBD. V_H_ Y99, Y100, and R100b have hydrophobic and π-π interactions with the RBD. The dashed box indicates potential space constraints for the glycine in the YYDRxG motif in CDR H3 as shown in Figure 2D.

### ADI-62113 binds a highly conserved site on SARS-CoV-2 RBD

To understand the basis of the broad reactivity of ADI-62113, we determined the crystal structure of ADI-62113 in complex with SARS-CoV-2 RBD at 2.6 Å resolution (Figure S1 and Table S1). ADI-62113 binds to the highly conserved CR3022 binding site (Yuan et al., 2020), but with an approach angle that can still block ACE2 receptor binding (Figure 1B), which in part explains its high neutralization potency against SARS-CoV-2. Of note, the approach angle and binding mode is very similar to another cross-neutralizing antibody, COVA1-16, that we reported previously, although the heavy chain variable domain of COVA1-16 is encoded by a different gene *IGHV1- 46* (Liu et al., 2020) (Figure S1A).

Further analysis of the antibody binding interface revealed that ADI-62113 CDR H3 dominates the interaction with SARS-CoV-2 RBD, contributing nearly 70% of the total buried surface area (BSA) on the RBD, as calculated by the PISA program (Krissinel & Henrick, 2007) (Figure 1C). The CDR H3 forms a β-strand like interaction with a SARS-CoV-2 RBD core β-strand through three main-chain to main-chain hydrogen bonds (Figure S1B). The aromatic ring of V_H_ Y99 interacts with aliphatic moiety of RBD K378 (Figure 1D). V_H_ Y100 forms a π-π interaction with the RBD peptide backbone of _382_VS_383_. V_H_ R94 interacts with the main-chain carbonyl of RBD G413, and V_H_ R97 with RBD D427 and F429 carbonyl oxygens. V_H_ Q101 forms equivalent hydrogen bond pairs with RBD Q414. The somatically mutated V_H_ R100b has multiple interactions with the RBD where its aliphatic moiety interacts with the aromatic ring of F377, and its guanidinium group forms three hydrogen bonds with the backbone of RBD _369_YNS_371_ and the partially negative dipole at the C-terminus of a short α-helix in the RBD, as also observed in COVA1-16 (Liu et al., 2020). The indented, negatively charged surface on the RBD in this region is highly suited for engagement with V_H_ R100b (Figure S2A). Of note, in the native spike trimer, this region is located in the interface between two adjacent RBDs, where this negative patch is engaged by R408 of a neighboring RBD in the “down” state, which may explain why arginine is naturally favored in ADI-62113 and COVA1-16 for interacting with this RBD site (Figure S2B). All of these epitope residues in the RBD are highly conserved across sarbecovirus sequences analyzed (Figure 1C and Figure S3), which supports the broad spectrum cross-reactivity observed for ADI-62113 against sarbecoviruses (Figure 1A).

### A YYDRxG motif shared between antibodies ADI-62113 and COVA1-16

Of note, we observed that CDR H3 of ADI-62113 exhibits near-identical interactions with the RBD relative to COVA1-16 despite differences in IGHV gene usage (Figure S1A-B). Interactions with other subsites seem have less impact on the binding mode as different interactions of RBD R408 with ADI-62113 and COVA1-16 light chains did not change the antibody approach angle or the binding mode of CDR H3. From comparison of these two structures, we identified that the centerpiece of the antibody paratopes, V_H 99_YYDRxG_100d_ hexapeptide, forms a conserved local structure for interaction with highly conserved residues in the RBD (Figure 2A).

**Figure 2.**
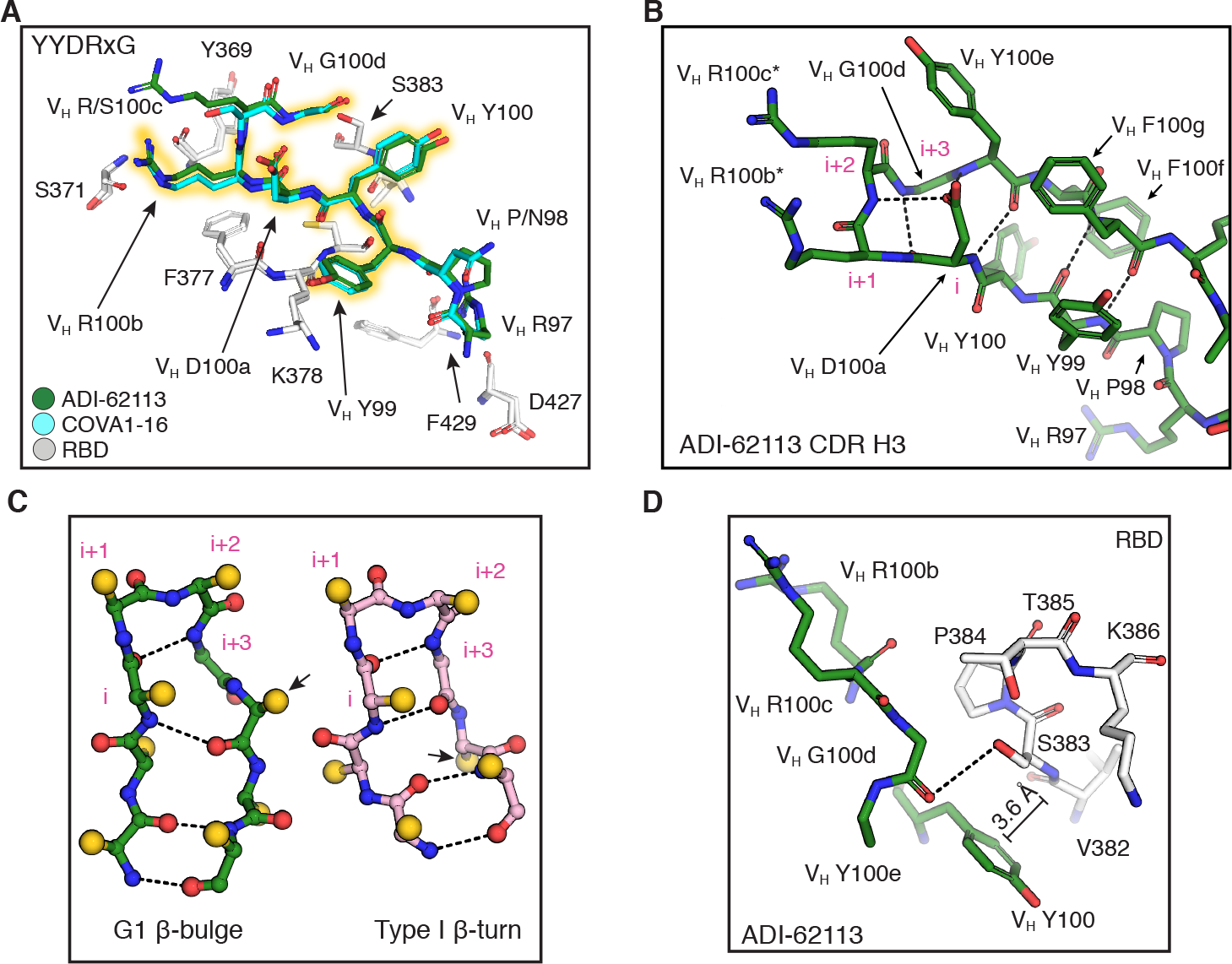
The YYDRxG motif is a recurring feature in CDR H3 for binding the RBD. Hydrogen bonds are shown as dashed lines. Key residues are shown in sticks. **A.** Comparison of CDR H3 interactions of ADI-62113 and COVA1-16 with the RBD. White, SARS-CoV-2 RBD; green, ADI- 62113; cyan, COVA1-16. The _99_YYDRxG_100d_ hexapeptide is highlighted to show the conserved structure in CDR H3 interacting with the RBD. **B.** The YYDRxG motif is located at the tip of CDR H3 and precedes a G1 β-bulge in the descending strand of the hairpin structure . Residues in the β-turn at the tip of CDR H3 are numbered i to i+3 (magenta). The V_H_ D100a carboxyl hydrogen bonds to backbone amides of V_H_ R100c (i+2) and V_H_ Y100e. *indicates somatically mutated residue. **C.** Backbone comparison of an inserted β-bulge versus a standard β-strand in a β-sheet. Schematic backbones show the β-harpin in ADI-62113 CDR H3 that contains a G1 β-bulge following the glycine residue (i+3) in the β-turn and comparison with a standard β-hairpin also with a type 1 β-turn at its tip (PDB ID: 4H5U). Amino-acid side chains are simplified as gold spheres. Arrows indicate the register change between the two motifs due to an additional residue in the β-hairpin in ADI-62113. **D.** Limited space between residue V_H_ G100d and RBD favors a glycine residue at this position in the β-bulge. A hydrogen bond is formed between the V_H_ G100d carbonyl oxygen and S383 hydroxyl in the RBD. A π-π interaction is formed between V_H_ Y100 and the peptide backbone of _382_VS_383_ of SARS-CoV-2 RBD.

A β-bulge is formed near the tip of CDR H3 (Figure 2B-C) after a type 1 β-turn (V_H 100a_DRRG_100d_ in this case) at its apex. An extra residue is inserted in the down strand of the β- hairpin at residue V_H_ Y100e and creates a G1 β-bulge, where the glycine residue in the bulge is in a left-handed α-helical conformation (Chan et al., 1993; Craveur et al., 2013; Richardson et al., 1978). The carbonyl oxygen of V_H_ D100a forms a typical hydrogen bond in type I β-turns to the amide of V_H_ G100d. The negatively charged side-chain carboxyl of V_H_ D100a is then able to hydrogen bond to the backbone amides of V_H_ R100c in the β-turn and V_H_ Y100e in the β-bulge, which may further stabilize the local structure of the β-bulge.

Glycine residues are often found in a β-bulge and almost exclusively required in a specific position of the G1 β-bulge due to the specific dihedral constraints at the position i+3 (Richardson et al., 1978) (n.b. normally first position in traditional G1 numbering but labeled as i+3 here to avoid confusion with β-turn numbering of the preceding turn). In this case, V_H_ G100d has a positive phi (φ) angle (φ=88°, ψ=11°), consistent with the glycine residue in a G1 β-bulge that alters the backbone conformation (Figure 2C) and enables a hydrogen bond to be made between its carbonyl oxygen and the hydroxyl of the highly conserved RBD S383 (Figure 2D). The presence of the additional residue in the G1 β-bulge motif alters the subsequent register of the β- sheet comprising the β-hairpin, thereby allowing for accommodation of an additional residue (Chan et al., 1993). The V_H_ Y100e side chain subsequently flips from a down configuration in the β-sheet to an up configuration (Figure 2C), thus opening up space for V_H_ Y100 to interact with the RBD (Figure 2D). The insertion of an extra residue into the β-sheet accentuates the typical right-handed twist of the β-sheet (Craveur et al., 2013), which may contribute to the structural stability and specificity of CDR H3 in its interaction with the RBD (Figure S1A). Moreover, the spatial constraint observed between the tip of CDR H3 of ADI-62113 and _382_VSPTL_386_ region in RBD may additionally drive the preference for a glycine in the β-bulge in ADI-62113 and COVA1-16 (Figure 2D and Figure S2C). Overall, this structural comparison of two cross-neutralizing antibodies, ADI- 62113 and COVA1-16, enabled identification of a unique pattern that includes the YYDRxG peptide where a register shift occurs in the β-hairpin through an insertion of a β-bulge. These features help promote twisting of β-sheet of the β-hairpin, contributing to potential stabilization of this particular conformation of the extended CDR H3. Importantly, the YYDRxG motif combines strong polar, hydrophobic, and π-π interactions with the CR3022 site on the RBD (Figure 2A), providing a structural basis for the potent and broad neutralization against SARS-CoV-2, VOCs, and other related viruses.

### Most YYDRxG motifs are encoded by *IGHD3-22* gene

Given these shared features were derived entirely from two cross-neutralizing antibodies (ADI- 62113 and COVA1-16), we sought to perform a computational pattern search to evaluate whether similar features might be present in any other antibodies with sequences deposited in public databases (Figure 3A). Of note, the CDR3 must be of sufficient length (here ≥18 residues by IMGT definition) to enable the YYDRxG hexapeptide to reach the conserved binding site on RBDs as observed for ADI-62113 and COVA1-16 (Figure S2D). Hence, length constraints in both N-terminal (≥5 aa) and C-terminal (≥7 aa) of the YYDRxG hexad were a requirement during the YYDRxG pattern search. Sequence homology to the YYDRxG hexapeptide was also used in the search since V_H_ Y99, V_H_ Y100, and V_H_ R100b all form hydrophobic interactions with the RBD, which may be possible with other residues containing hydrophobic moieties. In a search of over 205,000 antibody sequences, 153 antibodies with an YYDRxG pattern in their CDR H3 were identified (Tables S2-3). All sequence hits have the YYDRxG motif or close homologues in their CDR H3 (Figure S4A). Immunoglobulin gene analysis shows that the *IGHD3-22* gene is highly enriched. The D regions in CDR H3 in 88% of the YYDRxG antibodies are encoded by the *IGHD3- 22* (Figure 3B), which is the same diversity gene used in ADI-62113 and COVA1-16 (Figure S1A), compared to 8.5% antibodies in the overall search library (Figure S4B).

**Figure 3.**
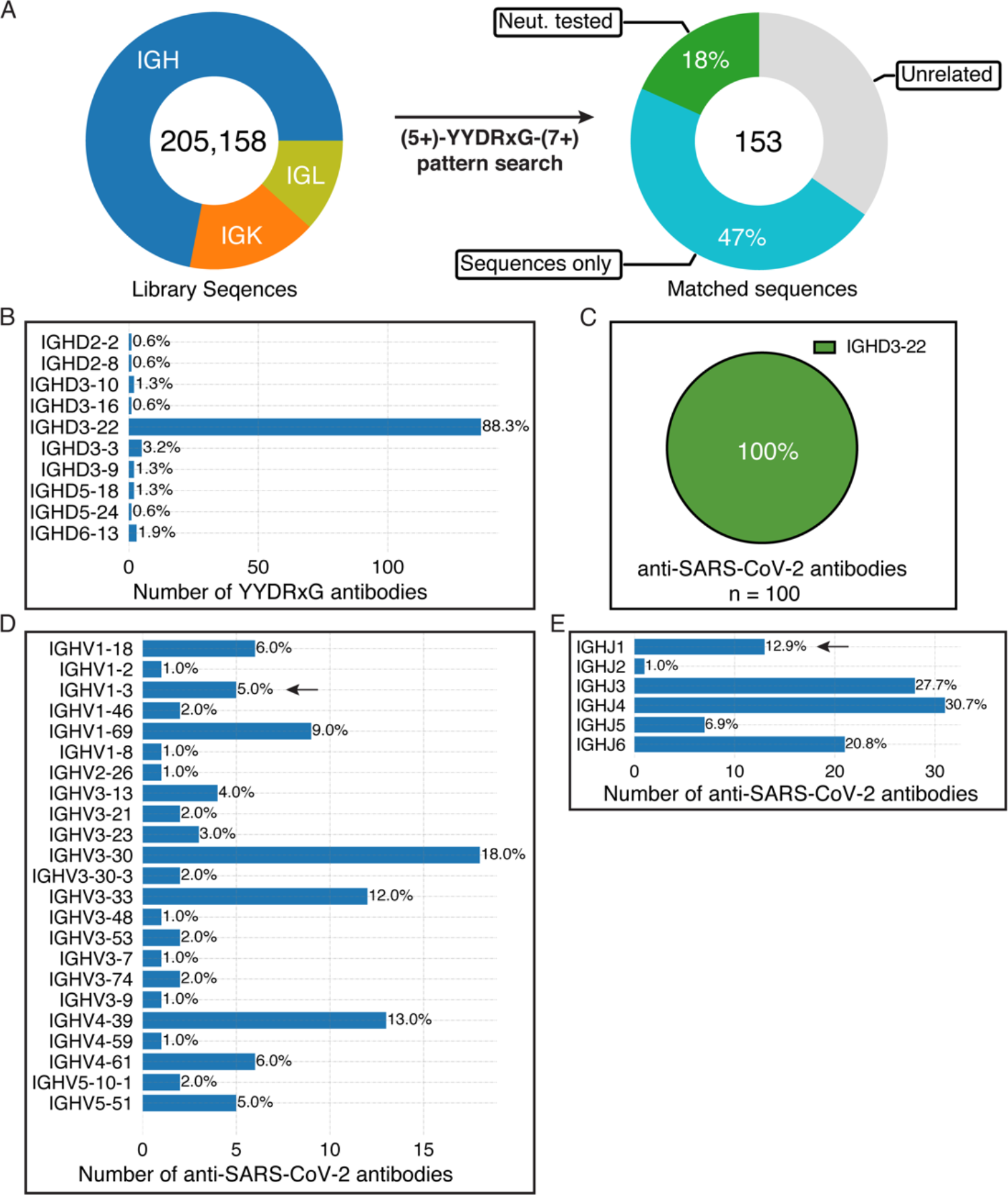
A YYDRxG pattern search identifies SARS-CoV-2 cross-neutralizing antibodies. **A.** Over 205,000 publicly available antibody sequences were retrieved from GenBank and supplementary files of previous publications reporting human anti-SARS-CoV-2 antibodies. The compiled antibody sequence library consists of 72% heavy chains, 16.4% kappa chains, and 11.7% lambda chains. CDR3 amino-acid sequences were used to search for the YYDRxG motif with five or more amino acid residues prior and seven or more amino acid posterior to the hexad motif. 153 heavy chain sequences were identified that contain a YYDRxG pattern and followed by data curation with literature review. 66% (n=100) are antibodies isolated from COVID-19 patients and mRNA vaccinees. 28 (18%) have been validated for SARS-CoV-2 neutralization (Neut. Tested), while 82 (47%) have not (sequences only). **B.** YYDRxG antibodies are mostly encoded by the *IGHD3-22* gene. All of the D genes used by YYDRxG antibodies were counted. Numbers at the right side of each bar indicate the percent of the total of identified YYDRxG antibodies. **C.** *IGHD3- 22* encodes all YYDRxG antibodies against SARS-CoV-2. All 110 sequences of anti-SARS-CoV- 2 antibodies were analyzed and counted. **D-E.** Heavy chain variable (D, IGHV) and joining (E, IGHJ) genes of YYDRxG antibodies isolated from COVID-19 patients or vaccinees. A diverse but limited set of variable and joining genes pairs with *IGHD3-22* in YYDRxG antibodies against SARS-CoV-2. *IGHV3-30, IGHV4-39, IGHV3-33,* and *IGHV1-69* are more frequently found than others. Arrows indicates the immunoglobulin genes that encode ADI-62113 in this study.

Of these 153 antibodies, 100 (65%) were isolated from different cohorts consisting of COVID-19 patients and mRNA vaccinees (Brouwer et al., 2020; E. C. Chen et al., 2021; Cho et al., 2021; Gaebler et al., 2021; Graham et al., 2021; Kreer et al., 2020; Robbiani et al., 2020; Sakharkar et al., 2021; Tong et al., 2021; Z. Wang, J. C. C. Lorenzi, et al., 2021; Z. Wang, F. Muecksch, et al., 2021; Z. Wang, F. Schmidt, et al., 2021). All of the D regions in these anti-SARS-CoV-2 antibodies are exclusively encoded by *IGHD3-22* (Figure 3C and Tables S3-4). In contrast, there does not appear to be a strict constraint on IGHV and IGHJ gene usage (Figure 3D-E), consistent with the different IGHV gene usage of ADI-62113 and COVA1-16. Diverse sets of IGHV and IGHJ genes are paired with *IGHD3-22* in these antibodies, which is distinct from, but overlap with, the YYDRxG antibodies identified in non-COVID cohorts (unrelated sources) (Figure S4C-D). Some IGHV genes are more often paired with *IGHD3-22* in these antibodies such as *IGHV3-30, IGHV4-39, IGHV3-33,* and *IGHV1-69,* although the current number of antibodies available for analysis (n=100 for anti-SARS-CoV-2 antibodies vs. n=53 for other antibodies) may limit conclusive determination of whether IGHV gene pairing preferences exist for this particular subset of SARS-CoV-2 antibodies.

### Somatic hypermutations, specific reading frame, and N addition to *IGHD3-22* are required

Antibody coding sequence is a product of immunoglobulin gene recombination, namely V(D)J recombination, non-templated addition (N addition), and somatic hypermutation during B cell development. The V(D)J junctional sequence encodes CDR H3 of an antibody. Three reading frames (RFs) are possible for the *IGHD3-22* due to random N additions, but the second RF is exclusively observed in these YYDRxG antibodies since the other two result in either early termination of translation (RF1) or a very different sequence (RF3) (Figure 4A). The situation differs from the *IGHD3-9* gene that is commonly used in broadly neutralizing anti-influenza stem antibodies, where two of the three RFs are used to encode bnAbs (Wu et al., 2018). Moreover, alignment of CDR H3 coding sequences of representative neutralizing antibodies against SARS-CoV-2 shows a high incidence of TàA/G or AàC transversions converting serine in the germline sequence to arginine (Figure 4B), supporting the hypothesis that somatic mutation from the germline serine to arginine residue in the YYDRxG motif (V_H_ R100b in ADI-62113 or COVA1-16) is critical for high affinity binding and neutralization (Liu et al., 2020). We also observe frequent somatic mutations adjacent to the serine codon (Figure 4A-B), which may be a lesion site created during antibody affinity maturation and serve as a prerequisite of converting S100b to R100b in germinal center as somatic hypermutation of A:T pairs requiring additional mutagenic process at neighboring sites (Di Noia & Neuberger, 2007; Franklin & Blanden, 2006). In addition to the S100b to R100b mutation in ADI-62113, mutation in a neighboring codon leads to somatically mutated R100c (Figure 2B and Figure 4B). While V_H_ R100b is critical for binding the RBD, V_H_ R100c has no interaction with RBD indicated by the paucity of side-chain electron density (Figure S2C). Thus, V_H_ R100c is not absolutely required since a serine at the same position does not impact binding, as observed in COVA1-16 (Figure S1B). N additions (N1 and N2) at both ends of *IGHD3-22* during V(D)J recombination seems important in determining the length of CDR H3 and the RF of *IGHD3- 22* that is critical in the positioning of YYDRxG in the CDR H3 β-hairpin. Overall, the requirement for a specific RF, site-specific somatic hypermutation, and relatively long N additions at both ends of *IGHD3-22* may contribute to the low frequency of YYDRxG motif or its homologues in the human antibody repertoire, which may in part explain the relatively rare occurrence of isolation of such cross-neutralizing antibodies in COVID-19 patients and vaccinees.

**Figure 4.**
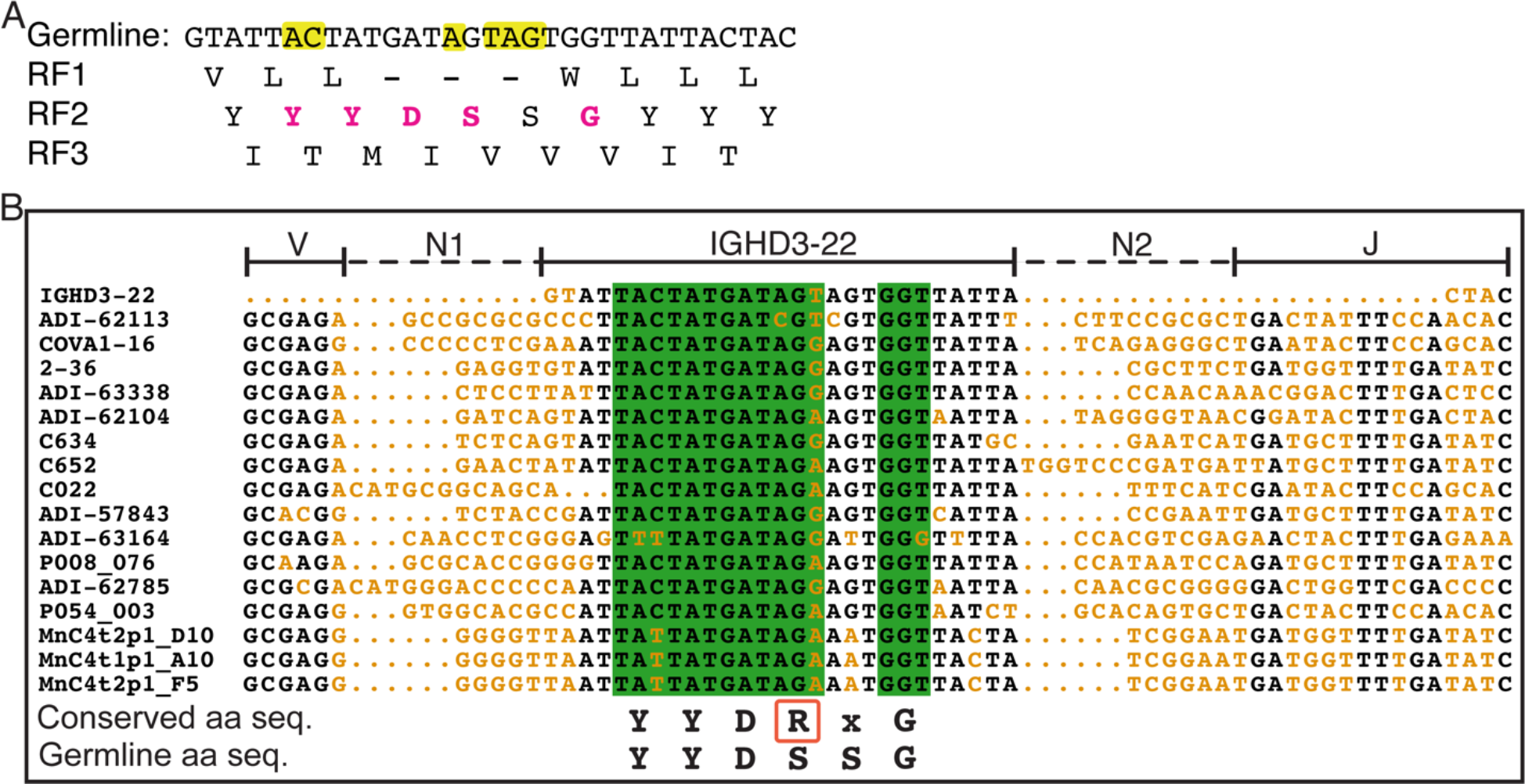
Reading frames (RFs) and sequence alignment of *IGHD3-22*. **A.** Three possible RFs of the *IGHD3-22* germline sequence were translated into amino-acid sequences. Only RF2 encodes the YYDRxG motif (magenta). Nucleotides highlighted in yellow in the germline DNA sequence indicate sites where somatic hypermutations are often found in YYDRxG neutralizing antibodies against SARS-CoV-2. **B.** Alignment of YYDRxG coding sequences in CDR H3. Conserved nucleotide sequences are in black, and variable sequences in orange. N additions (N1 and N2) as defined by IMGT junction analysis (Giudicelli & Lefranc, 2011) are indicated by dashed lines. *IGHD3-22,* IGHV(V), and IGHJ (J) regions are indicated by solid lines above the sequences. Sequences in a green background encode the YYDRxG key residues. Conserved YYDRxG sequence and corresponding germline amino-acid sequence (aa seq.) are shown below the sequence alignment panel for comparison. The arginine residue in the YYDRxG motif in a red box resulted from a high co-incidence of somatic mutations. Only the first 16 coding sequences are shown in the sequence alignment to represent 100 such antibodies against SARS-CoV-2.

### YYDRxG motif associates with potent and broad anti-SARS-CoV-2 antibodies

Among antibodies identified with a YYDRxG motif, 28 (18%) have been experimentally characterized via SARS-CoV-2 neutralization assays (Figure 3A). 25 out of 28 (89%) antibodies recognize SARS-CoV-2 RBD and 22 (79%) could effectively neutralize the virus as reported in previous publications (Figure 5A and Table S2) (Brouwer et al., 2020; Cho et al., 2021; Graham et al., 2021; Jette et al., 2021; Kreer et al., 2020; Sakharkar et al., 2021; Z. Wang, F. Muecksch, et al., 2021; Z. Wang, F. Schmidt, et al., 2021), although data for cross-reactivity to VOCs and other sarbecoviruses are incomplete. To determine whether the YYDRxG motif is associated with broad antigen recognition, we tested available ADI antibodies (Sakharkar et al., 2021) against a panel of sarbecovirus RBDs. We observe that all antibodies, except ADI-63219 and ADI-62969, strongly cross-react with multiple sarbecoviruses with apparent disassociation constants *K*_DApp_ ranging from 1.0 to 30.6 nM (Figure 5B and Figure S5). Despite the presence of the YYDRxG hexapeptide, ADI-62969 and ADI-63219 show weak affinity to all sarbecovirus RBDs tested including SARS-CoV-2. However, ADI-62969 and ADI-63219 have shorter or longer CDR H3 (19 or 25 residues by IMGT definition) compared to the others (average of 22 with a total range of 19- 26 residues) that may restrict the positioning of YYDRxG for binding to the RBD (Table S2-3). We further tested whether these antibodies cross-neutralize SARS-CoV-2 VOCs and SARS-CoV. Using pseudotyped viruses, these YYDRxG antibodies showed effective neutralization against Alpha, Beta, Gamma, and Delta VOCs, except again for ADI-63219 and ADI-62969 (Figure 5C and Figure S6). Consistent with the sarbecovirus binding data, five out of six cross-reactive antibodies neutralize SARS-CoV, albeit with variable potency ranging from 0.07 to >10 µg/ml. Combining with cross-neutralization data for four other antibodies (Graham et al., 2021; Jette et al., 2021; Liu et al., 2020; P. Wang, R. G. Casner, et al., 2021), nine (41%) of these 22 neutralizing antibodies could cross-neutralize SARS-CoV, albeit testing on another 11 (50%) of these antibodies against SARS-CoV may identify more such cross-neutralizing antibodies (Figure 5A).

**Figure 5.**
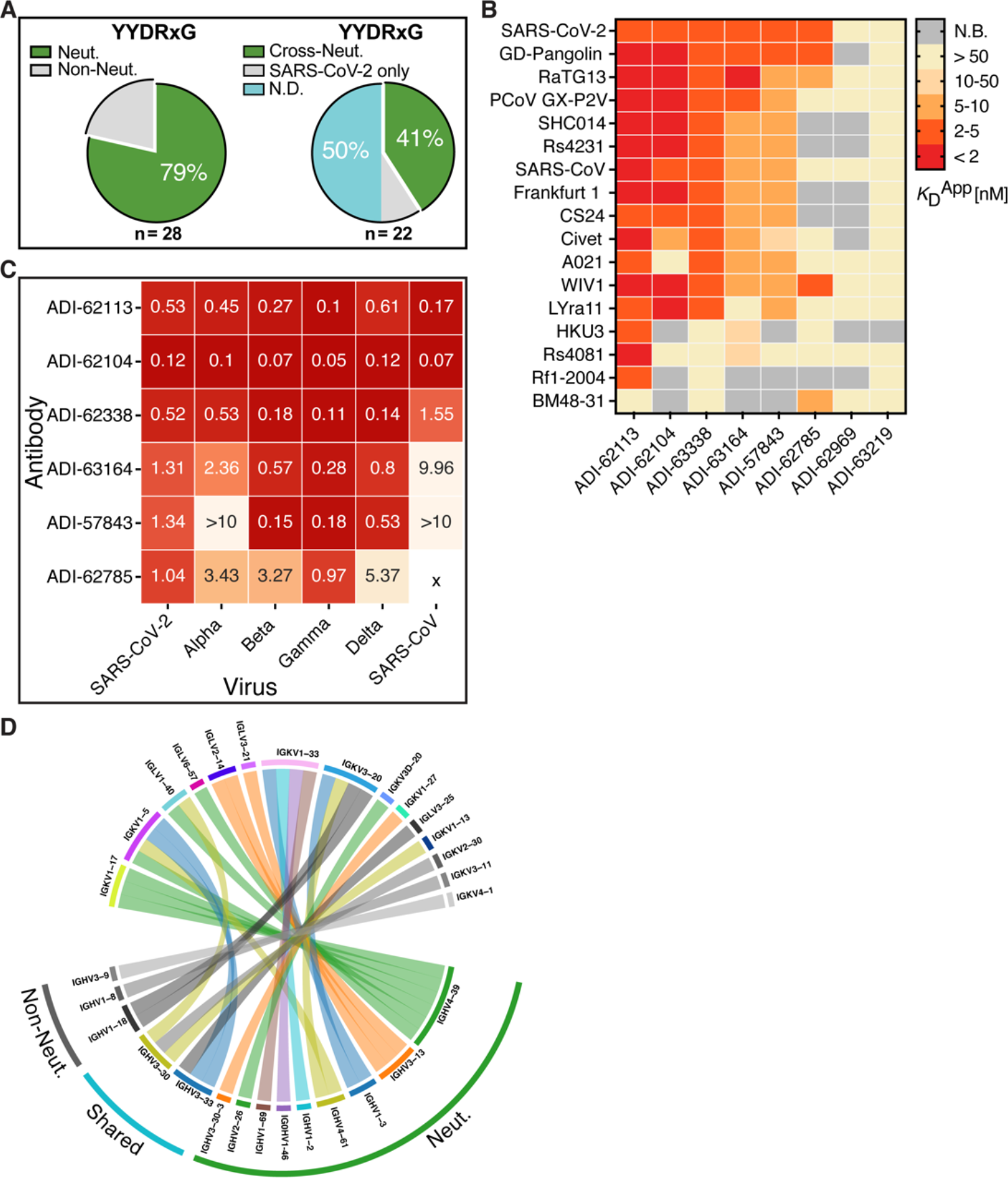
The YYDRxG motif is associated with potent and broad neutralization. **A.** Neutralization overview of YYDRxG antibodies against SARS-CoV-2. Antibodies with neutralization data available from previous publications or this study are included in this analysis. Within 28 YYDRxG antibodies, 22 (79%) exhibit neutralization (Neut.) against SARS-CoV-2 (left). Among these 22 antibodies, 9 (41%) cross-neutralize (Cross-Neut.) SARS-CoV, while 2 (9%) do not (SARS-CoV-2 only); another 11 (50%) have not been tested against SARS-CoV (N.D.) (right). **B.** Cross-reactivity of ADI antibodies with a YYDRxG motif across sarbecoviruses. ADI antibodies were titrated with sarbecovirus RBDs expressed on the yeast surface. Color bars indicated apparent binding affinities (*K*D^App^s) as indicated in the key. Red indicates strong binding, yellow indicates weak binding, and gray indicates no detectable binding (N.B.). Most antibodies are cross-reactive with many other sarbecovirus RBDs, except for ADI-63219 and ADI62969. ADI- 62113 showed the broadest spectrum of cross-reactivity to other sarbecoviruses. Titration curves used to determine binding affinities are shown in Figure S5. **C.** ADI antibodies neutralize SARS- CoV-2 VOCs and SARS-CoV. Neutralization was tested using a pseudovirus assay system. Neutralization potency, i.e. IC_50_, for each antibody against corresponding viruses are shown on a heatmap with red indicating potent neutralization. ADI-63219 and ADI-62969 showed no neutralization to SARS-CoV-2 (Figure S7) and are not included in this heatmap. X indicates no neutralization activity detected in the pseudovirus assay. **D.** Distinct combinations of heavy and light variable genes used by neutralizing and non-neutralizing antibodies. Circos plot showing combinations of heavy and light chain variable genes in each encoded antibody. Colored ribbons represent variable genes encoding neutralizing antibodies (Neut.) while gray indicate non- neutralizing antibodies (Non-Neut.). *IGHV3-30, IGHV3-33,* and *IGKV3-20* were found in both neutralizing and non-neutralizing antibodies. YYDRxG antibodies against SARS-CoV-2 with neutralization data available are included in this analysis (n=28). Antibody names and CDR H3 sequences of these antibodies are included in Table S2.

In addition, we analyzed antibodies from available neutralization data to see whether there was any preference in heavy chain or light chain variable gene segment usage paired with *IGHD3- 22* that was correlated with broad and/or cross-neutralization. We found several heavy and light chain V genes in neutralizing antibodies with no obvious preference in gene usage, although combinations of heavy and light variable genes in neutralizing antibodies are distinct from those in non-neutralizing antibodies (Figure 5D). *IGHV3-30, IGHV3-33,* and *IGKV3-20* were found to encode the heavy chain variable domain in both neutralizing and non-neutralizing antibodies. However, the small number of bnAbs that have been experimentally verified to date (n=22 neutralizing vs n=6 non-neutralizing antibodies) preclude statistically meaningful analysis.

Additional study of more YYDRxG antibodies will likely reveal how heavy and light chain genes and CDR H3 length might impact neutralization potency. Overall, our data suggest YYDRxG antibodies are broadly neutralizing antibodies for counteracting SARS-CoV-2 VOCs. Meanwhile, they are strongly associated with cross-species neutralization as eight of 10 tested antibodies exhibit cross-neutralization activity (Figure 5A and Table S2).

### YYDRxG pattern search identifies potent and broad antibodies against SARS-CoV-2 VOCs

Of the YYDRxG antibodies against SARS-CoV-2, more than two thirds have only sequence information available and have not been extensively evaluated or characterized (Figure 3A and Table S3). The strong correlation between the YYDRxG motif with experimentally characterized neutralizing antibodies suggested that these antibodies may broadly neutralize SARS-CoV-2 variants and potentially other sarbecoviruses, as observed in the antibodies that have been tested in the neutralization assay (Figure 5A and Table S2). Indeed, two representative antibodies with only sequence information available, i.e. MOD8_P2_IgG_B11-P1369 and PZF12_P2_IgG_F7- P1369, were selected from the computational search and showed broad neutralization against SARS-CoV-2, Alpha, Beta, Gamma, Delta, and SARS-CoV, although PZF12_P2_IgG_F7-P1369 has a lower potency against SARS-CoV. all the VOCs and SARS-CoV (Figure 6A). Further analysis of 80 YYDRxG antibodies isolated from 32 donors (Cho et al., 2021; Robbiani et al., 2020; Z. Wang, J. C. C. Lorenzi, et al., 2021; Z. Wang, F. Muecksch, et al., 2021; Z. Wang, F. Schmidt, et al., 2021) suggests that this type of antibody can be elicited in both infected and vaccinated patients, as well as in COVID-19 patients who received mRNA vaccines post-recovery (Figure 6B and Table S4). Collectively, these data strongly suggest the YYDRxG motif features in anti-SARS-CoV-2 antibodies that have potent and broad neutralization activity. The discovery of this YYDRxG pattern in antibodies against sarbecoviruses further support the notion that broad antibodies can be more frequently elicited in the human immune system than initially thought, which is consistent with recent publications where some vaccinee and patient sera exhibit detectable, although often diminished, neutralizing activity against SARS-CoV-2 VOCs and VOIs (Greaney et al., 2021; He et al., 2021; Jalkanen et al., 2021; J. Liu et al., 2021; Pegu et al., 2021; Z. Wang, F. Schmidt, et al., 2021; Wu et al., 2021; Yuan, Huang, et al., 2021).

**Figure 6.**
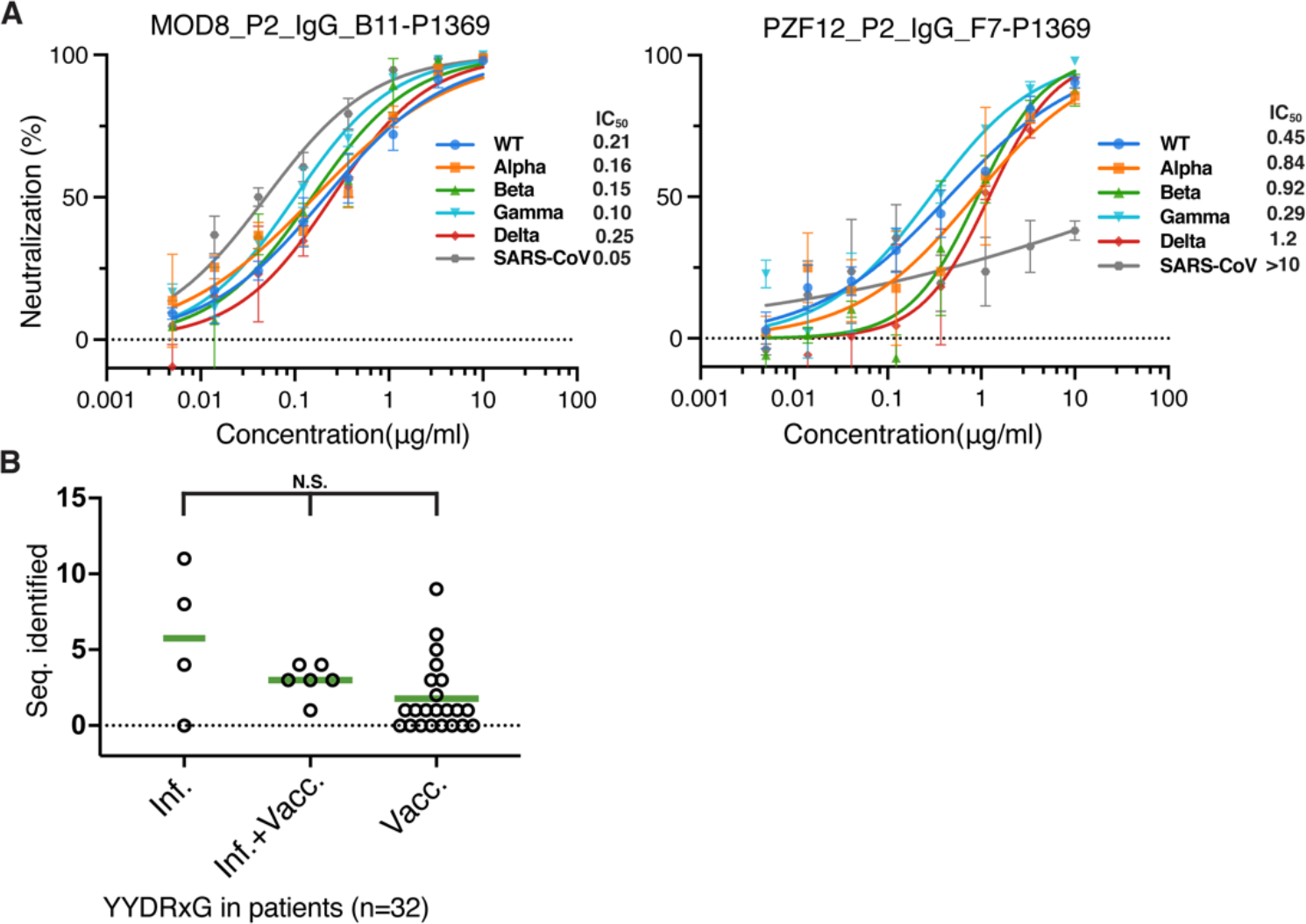
The YYDRxG pattern search identifies broad and potent antibodies to sarbecoviruses. **A.** Representative antibodies show potent neutralization. MOD8_P2_IgG_B11- P1369 and PZF12_P2_IgG_F7-P1369, were selected from the 83 sequences, which do not have any neutralization data available. The two antibodies were cloned, expressed, purified, and tested against SARS-CoV-2, VOCs and SARS-CoV in pseudovirus neutralization assay. Neutralization potency, i.e. IC_50_s are shown in the right of each panel. **B.** YYDRxG antibodies sequenced from COVID-19 and vaccinated patients. The number of YYDRxG antibody sequences identified in each patient’s antibodies (Seq. identified) were split into three groups according the medical history reported in their original publication (Cho et al., 2021; Gaebler et al., 2021; Robbiani et al., 2020; Z. Wang, J. C. C. Lorenzi, et al., 2021; Z. Wang, F. Muecksch, et al., 2021; Z. Wang, F. Schmidt, et al., 2021). All antibody sequences (n=17,536) sequenced from COVID-19 patients without vaccination (n=4), COVID-19 patients vaccinated post-recovery (n=6), and vaccinees without COVID-19 history (n=22) are included in this analysis. Non-parameter Kruskal-Wallis test was used to compare the number of YYDRxG antibodies isolated in each group. N.S., not significant (P>0.05). Sequence counts for each patient are included in Table S4.

## Discussion

Overall, we report here on structural characterization of a cross-neutralizing antibody, ADI-62113, that revealed a highly similar binding mode to the SARS-CoV-2 RBD that was first observed in COVA1-16 (Liu et al., 2020). We found that antibody binding to a very highly conserved site on the RBD appears to be determined primarily by the YYDRxG hexapeptide encoded by *IGHD3-22* in CDR H3. The extensive interaction with the conserved CR3022 site on the RBD and stabilization of the CDR H3 local structure by a β-bulge make the YYDRxG pattern a desirable feature for specific RBD recognition (Figure 2 and Figure S2A), which appears to be even more extensive than for the recurrent YYD sequences in HIV V2-apex targeting bnAbs (Andrabi et al., 2015; Gorman et al., 2016). Using the YYDRxG pattern search, we were able to identify and verify many more such antibodies as potent neutralizers of SARS-CoV-2, VOCs, and SARS-CoV.

Many RBS-targeting antibodies are highly potent for neutralizing specific SARS-CoV-2 viruses, but lack breadth of activity against emerging and circulating viruses (Andreano et al., 2021; R. E. Chen et al., 2021; Hoffmann et al., 2021; Starr, Greaney, et al., 2021; Tortorici et al., 2020; P. Wang, M. S. Nair, et al., 2021; Weisblum et al., 2020; Yuan, Huang, et al., 2021). The CR3022 site is highly conserved and therefore an important site for broad and cross-neutralizing antibodies (Liu et al., 2020; Yuan et al., 2020). Several potent antibodies, i.e. ADI-62113, COVA1- 16, C022, and S2X259, have now been structurally characterized that target the CR3022 site (Jette et al., 2021; Liu et al., 2020; Tortorici et al., 2021). Even although their epitopes do not overlap with ACE2, they compete with binding by the human ACE2 receptor, which may in part explain their relatively high potency in neutralizing both SARS-CoV-2 and VOCs.

High throughput methodologies especially single cell technologies and deep sequencing have advanced rapid antibody isolation to SARS-CoV-2 (Brouwer et al., 2020; Cao et al., 2020; Kreye et al., 2020; Robbiani et al., 2020; Rogers et al., 2020; Zost et al., 2020). Recently, repertoire sequencing has provided massive databases of antibody sequences (Briney et al., 2019; Soto et al., 2019). However, experimental characterization such as cloning, expression, kinetics measurement, epitope binning, and structure characterization are more time-consuming and sometimes resource limiting for studying the antibody response to SARS-CoV-2 infection and vaccination. From a search of public human antibody sequences, we identified 100 anti-SARS-CoV-2 antibodies containing a conserved YYDRxG motif exclusively encoded by *IGHD3-22* gene (Figure 3C and Table S2-3). The strong preference for one of the three possible reading frames of *IGHD3-22*, the apparent requirement for somatic hypermutation at a specific position in this region, local structural constraints, and the requirement for an extended length of CDR H3, make these antibodies less abundant compared to some other neutralizing antibodies targeting SARS-CoV-2 RBD. Our data and analysis show that antibodies with a YYDRxG pattern in their CDR H3 are more likely to target the highly conserved surface in sarbecovirus RBDs and to neutralize SARS-CoV-2 VOCs, emerging variants, and SARS-CoV, indicating a common convergent solution used by the human humoral immune system to counteract sarbecoviruses.

Finally, the high correlation between the presence of a YYDRxG pattern and broad neutralization activity support its use as a sequence signature of broadly and cross-neutralizing antibodies targeting this highly conserved site on the SARS-CoV-2 RBD as well as other related coronaviruses. Since antibodies containing a YYDRxG feature can be elicited by both natural infection and vaccination, interrogation of these signature sequences in serum can also serve as biomarkers to evaluate vaccine breadth and guide rational design of next-generation vaccines.

## MATERIALS AND METHODS

### Expression and purification of SARS-CoV-2 RBD

The receptor-binding domain (RBD) (residues 333-529) of the SARS-CoV-2 spike (S) protein (GenBank: QHD43416.1), was cloned into a customized pFastBac vector (Ekiert et al., 2011), and fused with an N-terminal gp67 signal peptide and C-terminal His_6_ tag (Yuan et al., 2020). Recombinant bacmids encoding each RBDs were generated using the Bac-to-Bac system (Thermo Fisher Scientific) followed by transfection into Sf9 cells using FuGENE HD (Promega) to produce baculoviruses for RBD expression. RBD protein was expressed in High Five cells (Thermo Fisher Scientific) with suspension culture shaking at 110 r.p.m. at 28 °C for 72 hours after the baculovirus transduction at an MOI of 5 to 10. Supernatant containing RBD protein was then concentrated using a 10 kDa MW cutoff Centramate cassette (Pall Corporation) followed by affinity chromatography using Ni-NTA resin (QIAGEN) and size exclusion chromatography using a HiLoad Superdex 200 pg column (Cytiva). The purified protein sample was buffer exchanged into 20 mM Tris-HCl pH 7.4 and 150 mM NaCl and concentrated for binding analysis and crystallographic studies.

### Expression and purification of antibodies

ADI-62113 IgG was produced in *S. cerevisiae* cultures, as previously described (Sakharkar et al., 2021). Briefly, yeast cultures were incubated at 30 °C and 80% relative humidity, with shaking at 650 rpm. Following 6 days of incubation, yeast cultures were cleared by centrifugation, and resultant supernatant containing IgG was purified by protein A chromatography. Bound IgGs were eluted from protein A resin using 200 mM acetic acid with 50 mM NaCl (pH 3.5) diluted into 1/8 [v/v] 2 M HEPES (pH 8.0) and subsequently buffer-exchanged into PBS (pH 7.0). Additional YYDRxG motif-containing monoclonal antibodies were produced with the same method for characterization by sarbecovirus RBD yeast surface display and neutralization assays. ADI- 62113 Fab fragments for structural studies were generated by papain digestion for 2 hours at 30 °C, and the reaction was terminated by addition of iodoacetamide. Fab fragments were purified over Protein A resin to remove cleaved Fc fragments and undigested IgG. The flowthrough was then purified over CaptureSelect™ IgG-CH1 affinity resin (ThermoFisher Scientific), and bound Fab fragments eluted with 200 mM acetic acid with 50 mM NaCl (pH 3.5) diluted into 1/8 [v/v] 2 M HEPES (pH 8.0) before buffer exchanging into PBS (pH 7.0). Expression plasmids encoding the heavy (HC) and light chains (LC) of Seq8 (MOD8_P2_IgG_B11-P1369) and Seq10 (PZF12_P2_IgG_F7-P1369) were transiently co-transfected into ExpiCHO cells at a ratio of 2:1 (HC:LC) using ExpiFectamine™ CHO Reagent (Thermo Fisher Scientific) according to the manufacturer’s instructions. The supernatant was collected at 10 days post-transfection. The IgG antibodies and Fabs were purified with a CaptureSelect™ CH1-XL Matrix column (Thermo Fisher Scientific) for affinity purification and a HiLoad Superdex 200 pg column (Cytiva) for size exclusion chromatography. The purified Fab protein samples were buffer exchanged into 20 mM Tris-HCl pH 7.4 and 150 mM NaCl and concentrated for crystallographic studies and the IgG proteins used for binding and neutralization assays.

### YYDRxG pattern search

Over 205,000 antibody sequences including heavy and light chains were retrieved from GenBank using Biopython program (Cock et al., 2009), supplemented with sequences reported in previous publications (Cho et al., 2021; Gaebler et al., 2021; Robbiani et al., 2020; Z. Wang, J. C. C. Lorenzi, et al., 2021; Z. Wang, F. Muecksch, et al., 2021; Z. Wang, F. Schmidt, et al., 2021), and then subjected to repertoire analysis using PyIR program (Soto et al., 2020) implemented with IgBLAST (Ye et al., 2013). CDR H3 amino-acid sequences from the compiled dataset were subjected to computational pattern search. Key residues in ADI-62113 and COVA1-16 interacting with SARS-CoV-2 RBD were analyzed using their crystal structures with SARS-CoV-2 RBD. YYDRxG and homologue sequences were used for the computational search. A length restriction with five or more amino acids prior and seven or more amino acids post YYDRxG key residues according to the structure analysis of ADI-62113 and COVA1-16 in complex with RBD was included in the search, which automatically yield a length restriction of >18 aa in CDR H3 that likely positions the YYDRxG motif to protrude towards the RBD surface. This constraint was thus defined along with the YYDRxG pattern to search for matches in compiled CDR H3 datasets. Genbank accession identifiers of the resultant YYDRxG hits were used for retrieving the full sequence record from Genbank. The compiled data were analyzed manually with sequence check and literature curation. Sequences of the antibodies against SARS-CoV-2 were included in further analysis.

### Sequence analysis and surface conservation

Sequences of sarbecoviruses were retrieved from GenBank using Biopython program, aligned using MUSCLE program (Edgar, 2004) built in European Bioinformatics Institute (EBI) web services (Madeira et al., 2019) and scored for surface conservation in ConSurf server (Ashkenazy et al., 2016; Landau et al., 2005). The conserved surface of SARS-CoV-2 RBD was visualized using the PyMOL program (Schrödinger, LLC). For visualization purpose, representative sarbecovirus RBD sequences from three clades were selected to compare with the conserved ADI-62113 epitope residues. BM48-31, YP_003858584; bat-Rf1-2004, ABD75323; PCoV_GX- P5L, QIA48632; PCoV_GX-P2V, QIQ54048; RaTG13, QHR63300; SARS-CoV-2, YP_009724390; A021-SARS-CoV-2, QWQ56573; bat-Rs4081, ATO98120; GD-pangolin, QLR06866; bat-LYRa11, AHX37558; civet010, AAU04649; SARS-CS24, ABF68959; bat-HKU3-1, AAY88866; SARS-CoV-Tor2, YP_009825051; Frankfurt 1, AAP33697; Rs4231, ATO98157; WIV1, AGZ48828; SHC014, QJE50589. Antibody sequences were also aligned using the same method and visualized using ESPript 3.0 server (Robert & Gouet, 2014).

### Crystallization and X-ray structure determination

The ADI-62113 Fab was mixed with SARS-CoV-2 RBD in an equimolar ratio and incubated overnight at 4°C. 384 conditions of the JCSG Core Suite (Qiagen) were used for setting-up trays for screening at a concentration of 11.9 mg/ml on our robotic CrystalMation system (Rigaku) at Scripps Research. Crystallization trials were set-up by the vapor diffusion method in sitting drops containing 0.1 μl of protein complex and 0.1 μl of reservoir solution. Crystals appeared on day 2, were harvested on day 14, and flash cooled and stored in liquid nitrogen until data collection. Diffraction data were collected at cryogenic temperature (100 K) at the Stanford Synchrotron Radiation Lightsource (SSRL) on Scripps/Stanford beamline 12-1 with a beam wavelength of 0.97946 Å, and processed with HKL2000 (Otwinowski & Minor, 1997). Diffraction data were collected from crystals grown in a drop containing 0.08 M sodium acetate pH 4.6, 0.16 M ammonium sulfate, 20% (w/v) polyethylene glycol 4000, and 20% (v/v) glycerol. The X-ray structure was solved by molecular replacement (MR) using PHASER (McCoy et al., 2007) with MR models for the RBD and Fab from PDB 7JMW (Liu et al., 2020). Iterative model building and refinement were carried out in COOT (Emsley & Cowtan, 2004) and PHENIX (Acta Crystallographica Section D: Biological CrystallographyAdams et al., 2010), respectively. Epitope and paratope residues, as well as their interactions, were identified with the PISA program (Krissinel & Henrick, 2007) using calculated buried surface area (BSA >0 Å^2^) as the criterion.

### Binding to sarbecovirus RBDs via yeast surface display

To assess binding breadth, IgGs and recombinant human Fc-conjugated hACE2 (Sino Biological, 10108-H02H) were tested against the panel of 17 sarbecovirus RBDs expressed by yeast display as previously described (Rappazzo et al., 2021). Briefly, EBY100 yeast were transformed with a plasmid encoding the yeast mating protein Aga2p linked to sarbecovirus RBD on the C terminus. To induce RBD expression, 0.5 OD_600_/ml of yeast were transferred to SGCAA media and cultured at 20°C for 16-20 hours with 180 rpm shaking. Next, IgGs and hACE2 were titrated via 3-fold serial dilutions from 100 nM to 0.5 pM. RBD-expressing cells were aliquoted into 96-well plates and incubated with 100 µl of 100 nM IgG for 30 minutes on ice. Next, cells were washed twice with PBSF (1X PBS, 0.1% BSA) before secondary detection with 1:100 dilutions of APC-conjugated mouse anti-hemagglutinin tag (HA).11 antibody (BioLegend, 901524), PE-conjugated goat anti-human IgG polyclonal antibodies (Southern Biotech, 2040-09), and propidium iodide (Invitrogen, P1304MP) for 20 minutes on ice. Cells were washed twice with PBSF before analyzing via flow cytometry on a BD FACS Canto II (BD Biosciences). Binding curves were fitted with four-parameter non-linear regression analysis to calculate the apparent equilibrium binding constant (*K*D^App^) in GraphPad Prism 9. Points exhibiting hook effects at higher concentrations were excluded from analysis.

### Pseudovirus production and neutralization assays

Plasmids encoding SARS-CoV, SARS-CoV-2, or other variants of the spike protein, with the ER retrieval signal removed were co-transfected with MLV-gag/pol and MLV-CMV-Luciferase plasmids into HEK293T cells to generate pseudoviruses. Lipofectamine 2000 (Thermo Fisher Scientific, 11668019) was used according to the manufacturer’s instructions. The cell culture supernatants containing S-pseudotyped MLV virions were collected at 48 hours post transfection, filtered through a 0.22 μm membrane and stored at -80°C until use. Lentivirus transduced Hela cells expressing hACE2 (GenBank: BAB40370.1) were enriched by fluorescence-activated cell sorting (FACS) using biotinylated SARS-CoV-2 RBD conjugated with streptavidin-Alexa Fluor 647 (Thermo, S32357). Stable cell lines with consistent and high hACE2 expression levels were established as HeLa-hACE2 and used in the pseudovirus neutralization assay. Monoclonal antibody IgGs were serially diluted with DMEM medium supplemented with 10% heat-inactivated FBS, 1% Q-max, and 1% P/S. 25 μl of 3-fold serial dilutions were incubated with 25 μl of pseudotyped viruses at 37°C for 1 hour in 96-well half-well plate (Corning, 3688). Right before the end of the incubation, HeLa-hACE2 cells were suspended with culture medium at a concentration of 2 x 10^5^/ml. The DEAE-dextran (Sigma, 93556-1G) was added to the cell solutions at 20 μg/m for enhanced infectivity. 50 μl of the cell solution was distributed into each well. The supernatant was removed 48 hours post incubation at 37°C, and the neutralization efficiency was calculated by measuring the luciferase levels in the HeLa-hACE2 cells. Cells were washed and lysed in luciferase lysis buffer (25 mM Gly-Gly pH 7.8, 15 mM MgSO_4_, 4 mM EGTA, 1% V/V Triton X-100) at room temperature for 10-20 mins. After addition of Bright-Glo (Promega, PR-E2620) according to the manufacturer’s instruction, luminescence signal was measured in duplicate. At least two biological replicates were performed for neutralization assays with SARS-CoV-2, SARS- CoV-2 variants of concern, and SARS-CoV. The IgG half-maximal inhibitory concentration (IC_50_) values were calculated using “One Site - Fit LogIC50” regression in GraphPad Prism 9.

## Acknowledgements

We thank Henry Tien for technical support with the crystallization robot, Jeanne Matteson and Yuanzi Hua for contribution to mammalian cell culture, Wenli Yu for insect cell culture, Seiya Kitamura for useful discussions, and Robyn Stanfield for assistance in data collection. We are grateful to the staff of Stanford Synchrotron Radiation Lightsource (SSRL) Beamline 12-1 for assistance. This work was supported by Adagio Therapeutics and by the Bill and Melinda Gates Foundation INV-004923 (I.A.W., D.R.B.). This research used resources of the SSRL, SLAC National Accelerator Laboratory, which is supported by the U.S. Department of Energy, Office of Science, Office of Basic Energy Sciences under Contract No. DE-AC02–76SF00515. The SSRL Structural Molecular Biology Program is supported by the DOE Office of Biological and Environmental Research, and by the National Institutes of Health, National Institute of General Medical Sciences (including P41GM103393). Extraordinary facility operations were supported in part by the DOE Office of Science through the National Virtual Biotechnology Laboratory, a consortium of DOE national laboratories focused on the response to COVID-19, with funding provided by the Coronavirus CARES Act.

## Author Contributions

H.L., C.I.K, L.M.W., and I.A.W. conceived and designed the study. H.L. and C.I.K. expressed and purified the proteins for crystallization and the binding and neutralization assays. H.L. crystallized the antibody-antigen complexes and determined the crystal structures. M.Y. helped with X-ray diffraction data collection. H.L. and I.A.W. analyzed the structural data. H.L performed the antibody data mining and sequence analysis. C.I.K. and L.M.W. provided binding antibody breadth data. G.S., R.A., and D.R.B. provided neutralization data. H.L. and I.A.W wrote the paper and all authors reviewed and/or edited the paper.

## Competing Interest Statement

C.I.K. and L.M.W. are employees of Adimab, LLC and hold shares in Adimab, LLC. L.M.W. is an employee of Adagio Therapeutics Inc. and holds shares in Adagio Therapeutics Inc. L.M.W. is an inventor on a patent describing the ADI anti-SARS-CoV-2 antibodies.

## SUPPLEMENTAL FIGURE AND FIGURE LEGENDS

**Figure S1.**
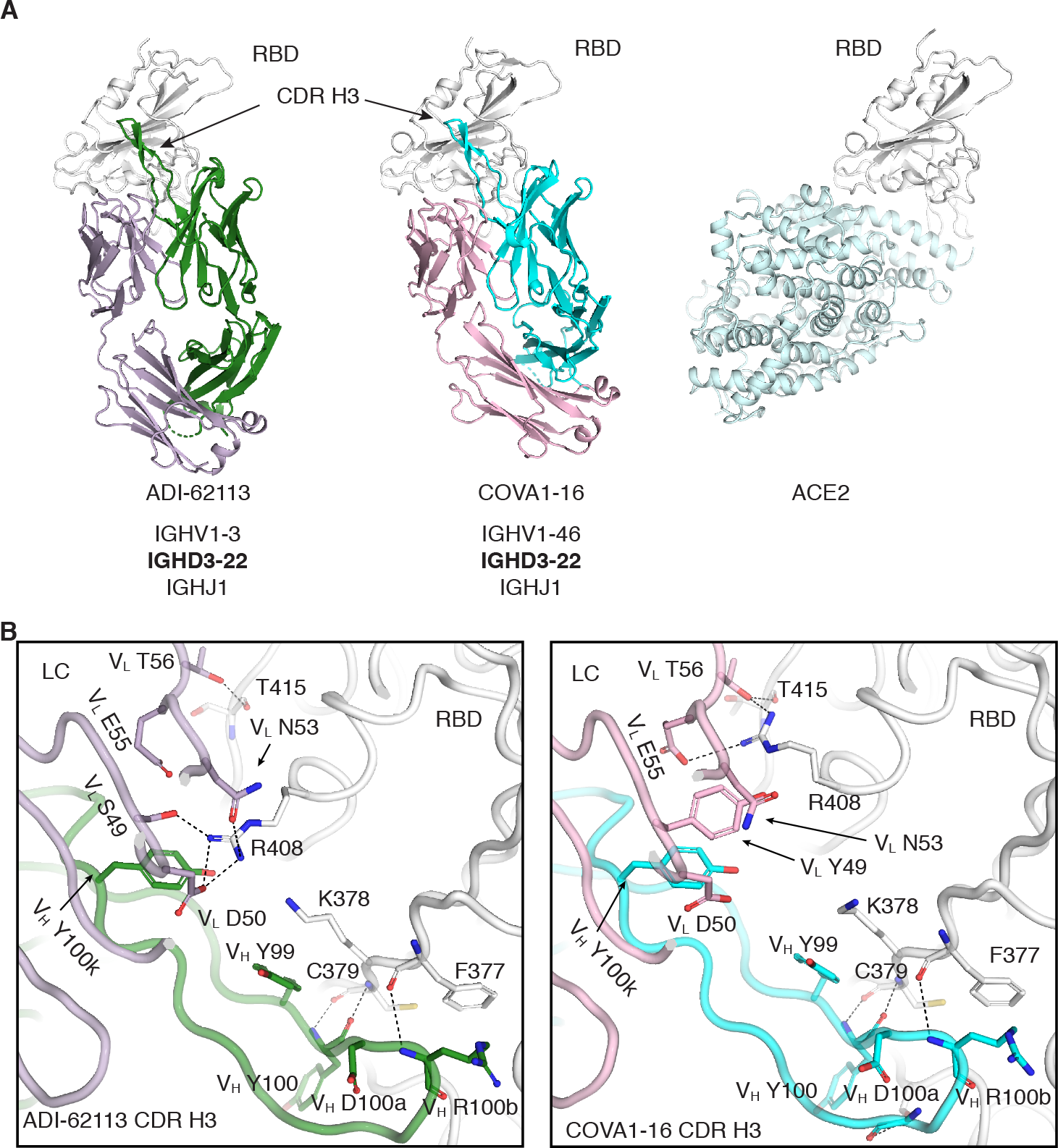
Structural comparison of ADI-62113 and COVA1-16. **A.** A similar binding mode to SARS-CoV-2 RBD was found for ADI-62113 and COVA1-16. Structures are shown in ribbon representation. Both antibodies compete with ACE2 binding as shown in the far right even although their epitopes do not directly overlap with ACE2. SARS-CoV-2 RBD is shown in white. The same perspective view was used for all structures. The β-hairpin in CDR H3 is more twisted compared to the strands in the β-sheet in the core region of the Fab. Heavy chain immunoglobulin genes are shown under each antibody for comparison. Both antibodies use the *IGHD3-22* gene encoding the YYDRxG motif. **B.** Comparison between ADI-62113 and COVA1-16 Fab binding to SARS-CoV-2 RBD. ADI-62113 adopts a similar binding mode to COVA1-16. A long CDR H3 containing the YYDRxG motif dominates the binding to SARS-CoV-2. Some differences in detailed interactions are seen due to sequence variation between the different antibodies, e.g. interaction with R408 of the RBD. The structures are shown in backbone tube representation with the side chains of residues involved in antibody-antigen interaction in sticks.

**Figure S2.**
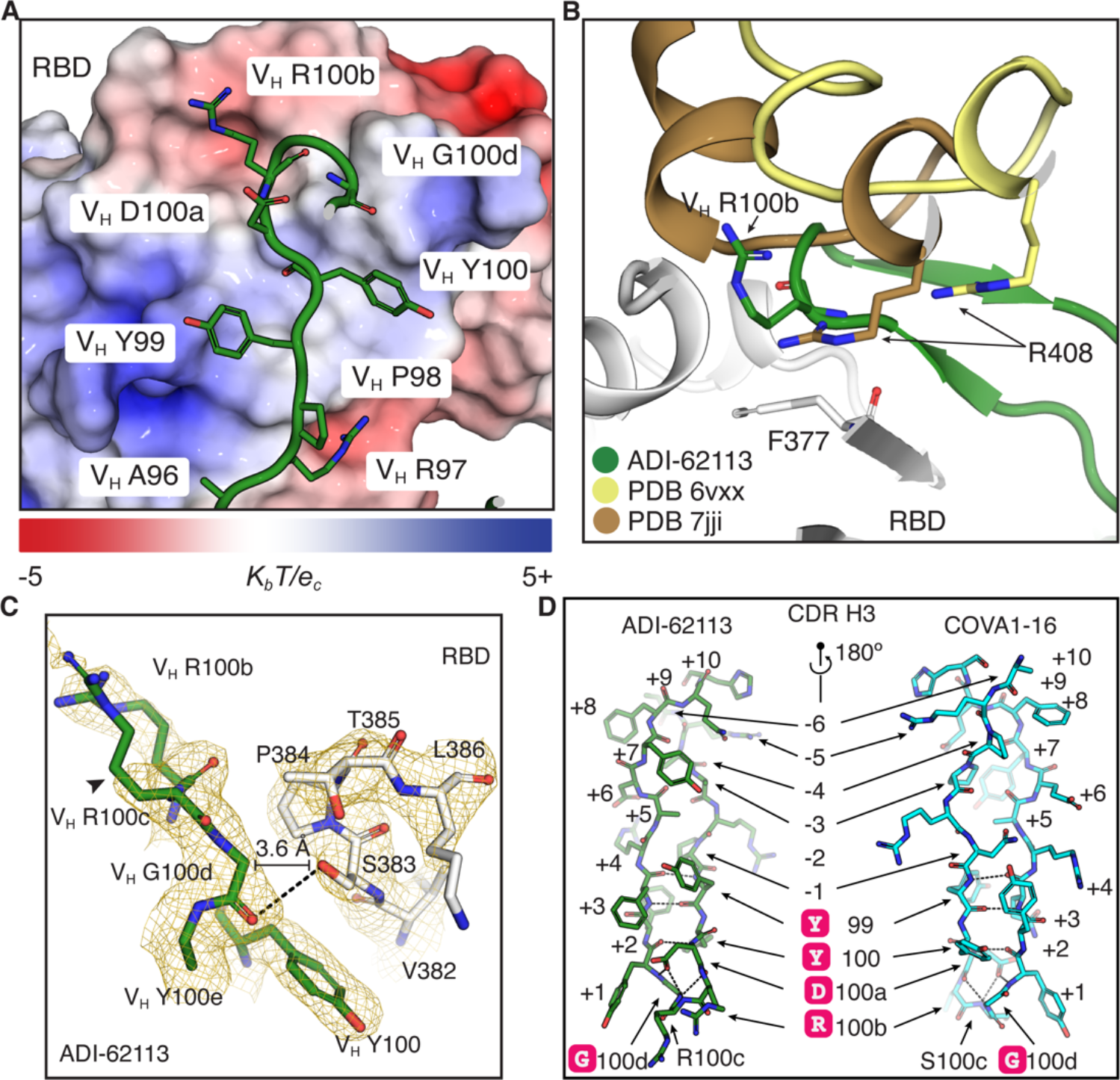
YYDRxG motif analysis. **A.** CDR H3 of ADI-62113 bound to SARS-CoV-2 RBD surface. SARS-CoV-2 RBD is shown in an electrostatic surface representation. Red indicates a negatively charged surface and blue indicates a positively charged surface. The bar below the panel shows the electrostatic scale. CDR H3 is shown in a backbone tube representation with side chains in sticks. V_H_ R100b binds to a recessed surface on the RBD with an overall negative charge. **B.** The conserved surface where V_H_ R100b binds is also part of the interface between spike protomers when RBD is in the down conformation. Intra-protomer interaction among RBDs within a spike is variable due to slight differences among RBD conformations in different structures. Spikes with all RBD in “down” state are shown using two PDB IDs: 6VXX and 7JJI. R408 from the neighboring RBD can bind to the same site where V_H_ R100b in ADI-62113 and COVA1-16 binds. Interaction between an arginine and F377 also seems to be common for both intra-protomer binding and antibody recognition at this site. **C.** Electron density around ADI-62113 V_H_ R100c. The electron density map is contoured at σ=1.0 using a 2mFo-DFc map calculated from the structure factors using PHENIX program (Acta Crystallographica Section D: Biological CrystallographyAdams et al., 2010). The V_H_ R100c has poor to no side-chain density likely due to high mobility (indicated by an arrowhead). **D.** Comparison of the shared features of YYDRxG in ADI-62113 and COVA1-16. The hydrogen bonding observed with the YYDRxG hexapeptide are almost identical, indicating a conserved structure of the two CDR H3s. Sequences outside of YYDRxG are highly variable and two proline substitutions (at positions -3 and -4) have no impact on the YYDRxG conformation. The distance between YYDRxG to the first and last residue of CDR H3 seems important to position the hexapeptide at the tip of CDR H3 for RBD binding. A minimum of 5+ aa prior and 7+aa post YYDRxG hexapeptide should allow formation of a similar CDR H3 structure.

**Figure S3.**
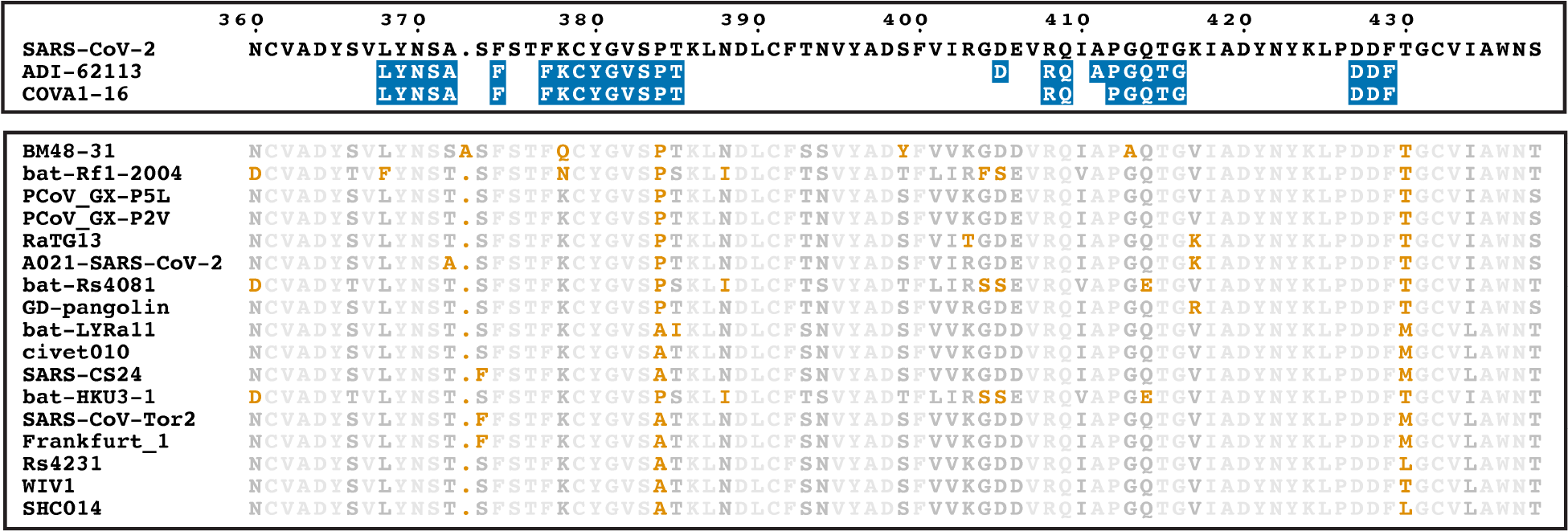
Conserved epitope residues of YYDRxG antibodies. Epitope residues of ADI- 62113 and COVA1-16, two YYDRxG antibodies, are highly conserved across sarbecoviruses. Epitope residues are highlighted on the sequence of SARS-CoV-2 RBD in the upper panel. Epitope residues are defined by having the buried surface area (BSA) by the indicated antibody with BSA >0 Å^2^ as the criterion. Other representative sarbecovirus sequences are aligned in the lower panel. Conserved residues are shown in brown, variable in brown. RBD amino-acid sequences were retrieved from GenBank database with the following IDs: BM48-31, YP_003858584; bat-Rf1-2004, ABD75323; PCoV_GX-P5L, QIA48632; PCoV_GX-P2V, QIQ54048; RaTG13, QHR63300; SARS-CoV-2, YP_009724390; SARS-A021, QWQ56573; bat- Rs4081, ATO98120; PCOV-GD, QLR06866; bat-LYRa11, AHX37558; civet010, AAU04649; SARS-CS24, ABF68959; bat-HKU3-1, AAY88866; SARS-CoV-Tor2, YP_009825051; Frankfurt 1, AAP33697; Rs4231, ATO98157; WIV1, AGZ48828; SHC014, QJE50589.

**Figure S4.**
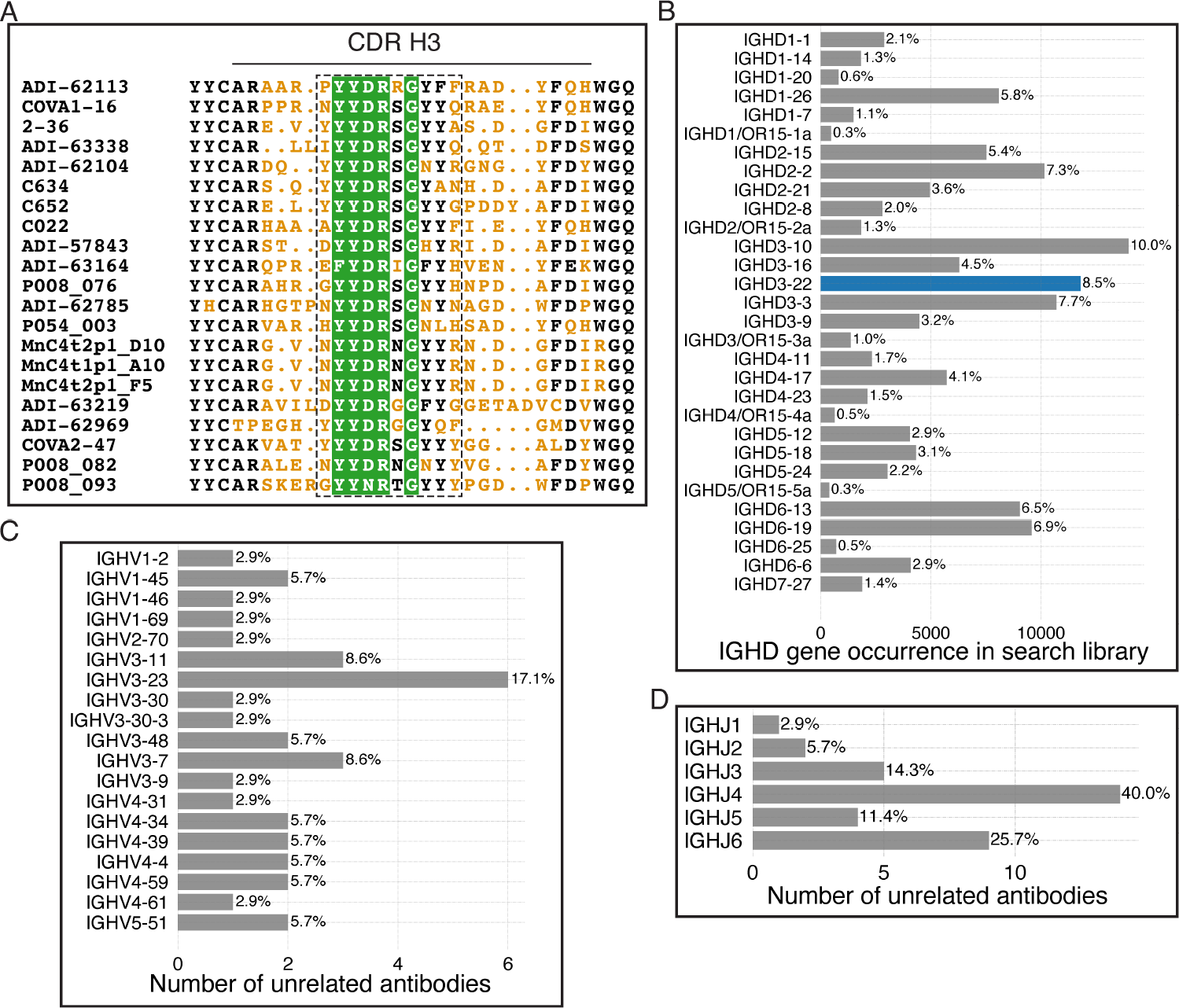
YYDRxG antibodies identified from the publicly available sequences. **A.** CDR H3 sequence alignment of representative YYDRxG antibodies against SARS-CoV-2. Conserved residues are shown in black and white, and variable residues in orange. Green shades highlight key residues in the YYDRxG motif. CDR H3 sequences for all antibodies are included in Table S2 (tested against SARS-CoV-2) and Table S3 (sequence only). CDR H3 regions (IMGT definition) are indicated by the solid line above the sequences. Sequences encoded by *IGHD3-22* are boxed with a dashed line. **B.** Immunoglobulin diversity (IGHD) genes in the computational search library compiled with publicly available antibody sequences are described in Methods section. IGHD genes are diversely distributed in the overall library. No single IGHD gene is dominant in the antibody sequences, although some have higher propensity than others. *IGHD3-22* encoding YYDRxG antibodies is highlighted. **C-D.** Immunoglobulin variable (IGHV) and joining (IGHJ) genes in YYDRxG antibodies isolated from non-COVID-19 cohorts. The diverse sets of IGHV and IGHJ genes are distinct from, but overlap with, those observed in anti-SARS-CoV2 antibodies.

**Figure S5.**
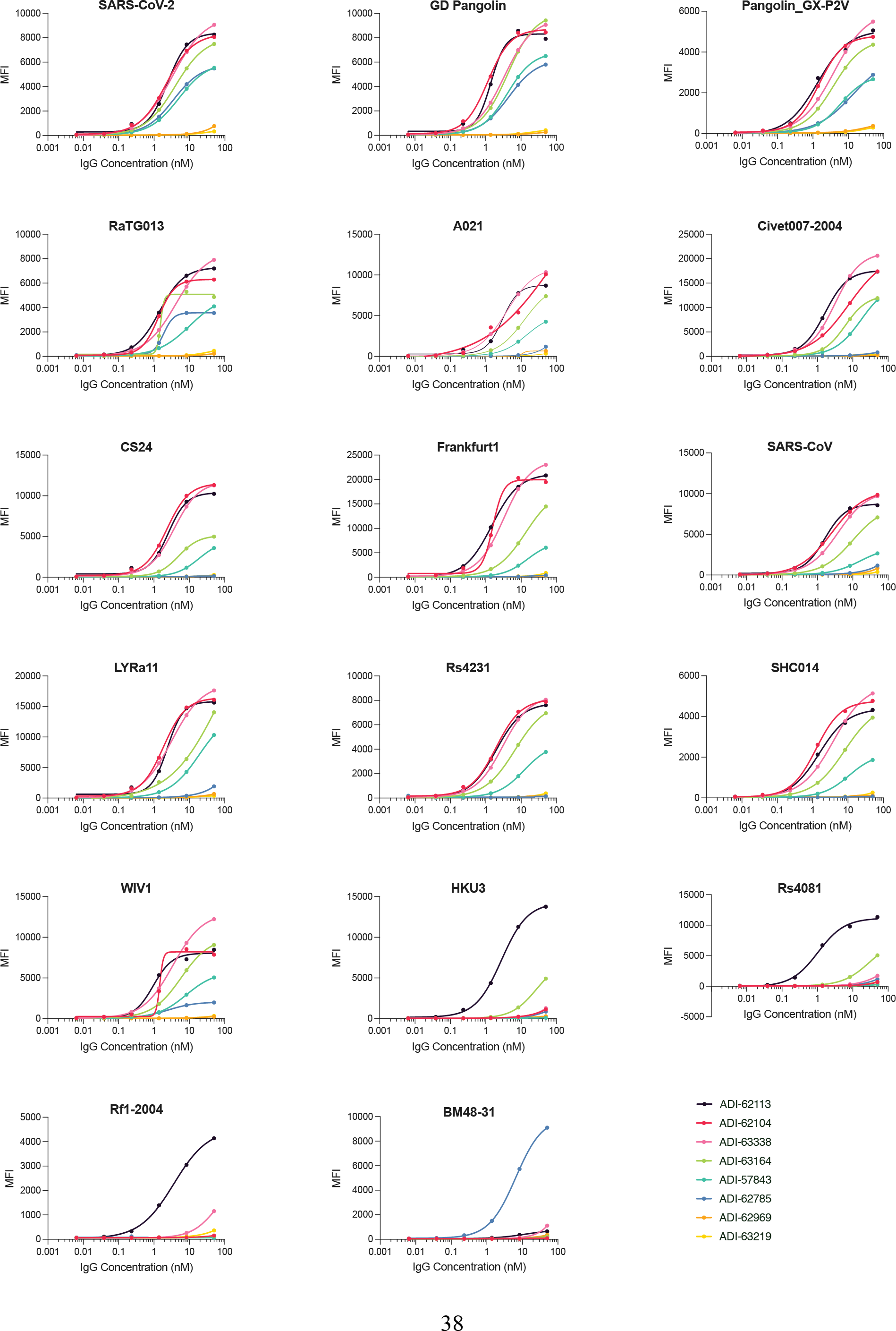
Cross-reactivity of YYDRxG antibodies with sarbecoviruses. ADI antibodies were used for kinetic binding assay with sarbecovirus RBDs displayed on the yeast cell surface. Binding affinities are plotted as a heatmap in Figure 5B. Most antibodies show potent binding to multiple sarbecoviruses, except for ADI-62969 and ADI-63219. The CDR H3 length of these two antibodies seems to be consistent with diminished binding to antigen.

**Figure S6.**
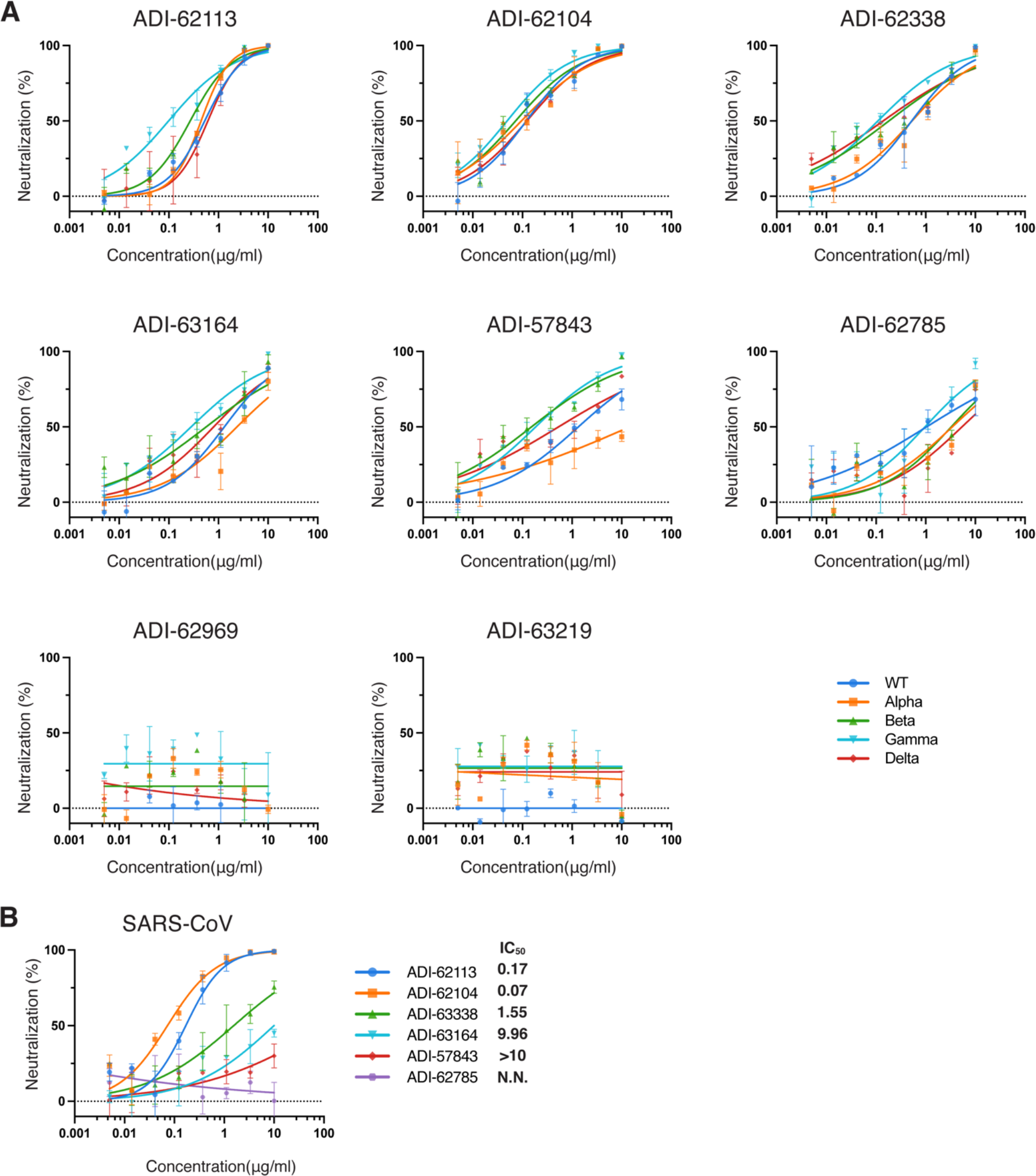
YYDRxG antibodies neutralize SARS-CoV-2 VOCs and SARS-CoV. ADI antibodies were tested in a neutralization assay against SARS-CoV-2 wild type (WT), VOCs, and SARS-CoV pseudoviruses. Neutralization potencies, IC_50_s, are shown in Figure 5C. Consistent with the binding kinetics assay, most antibodies broadly neutralize SARS-CoV-2 variants (A) and SARS- CoV (B).

## SUPPLEMENTAL TABLE AND TABLE LEGENDS

**Table S1.**
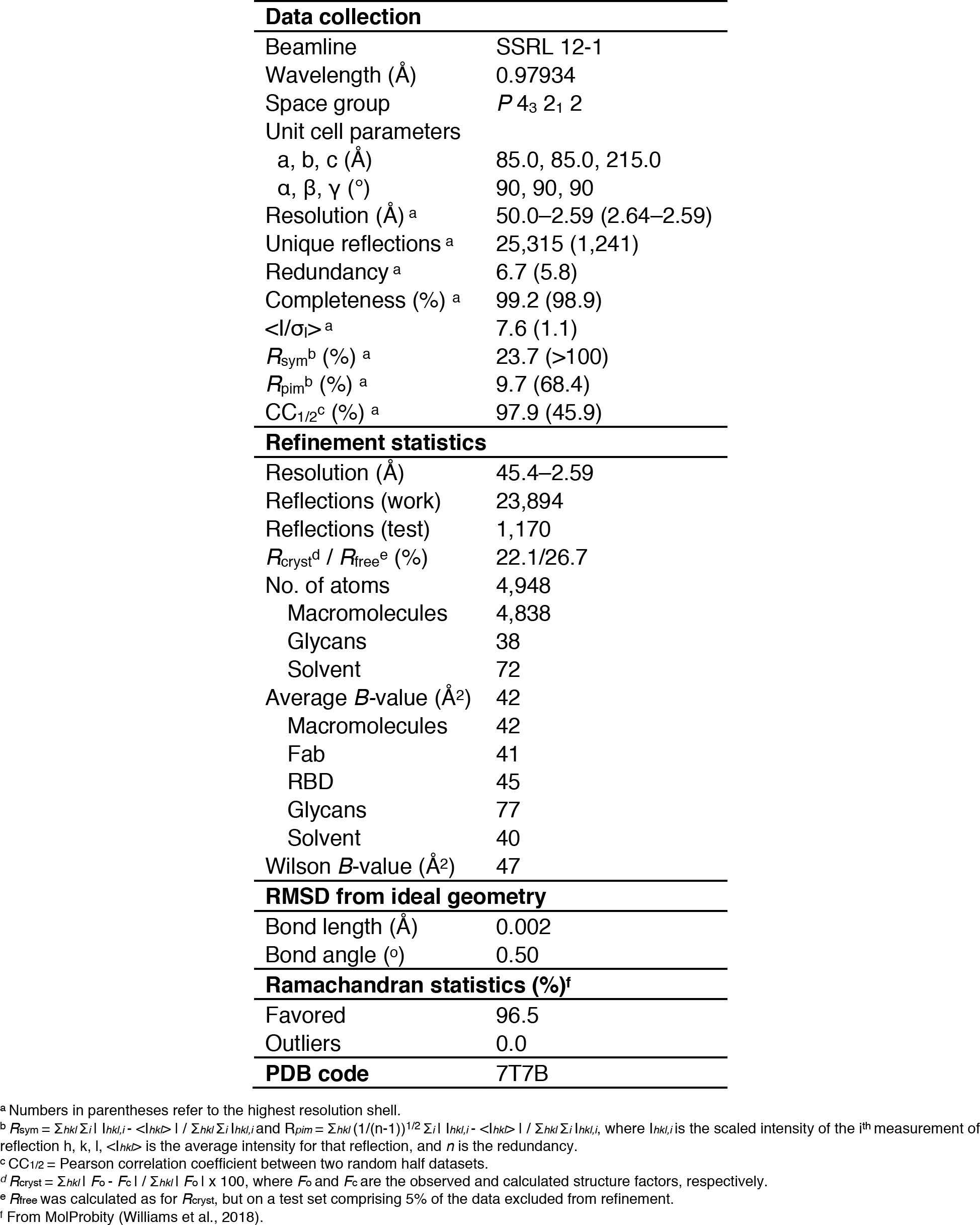
Crystallographic data collection and refinement statistics

**Table S2.**
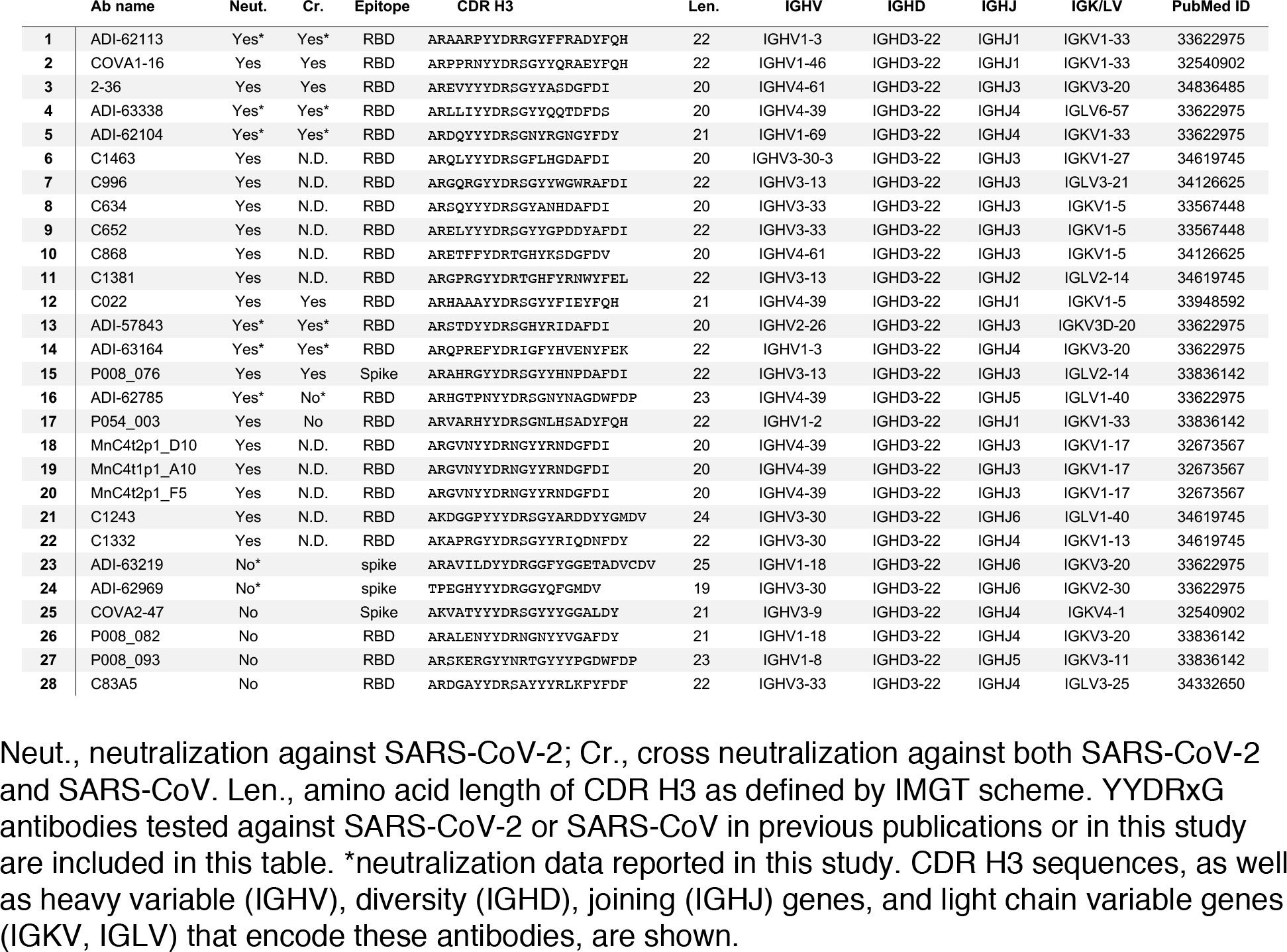
YYDRxG antibodies against SARS-CoV-2 with available neutralization data

**Table S3.**
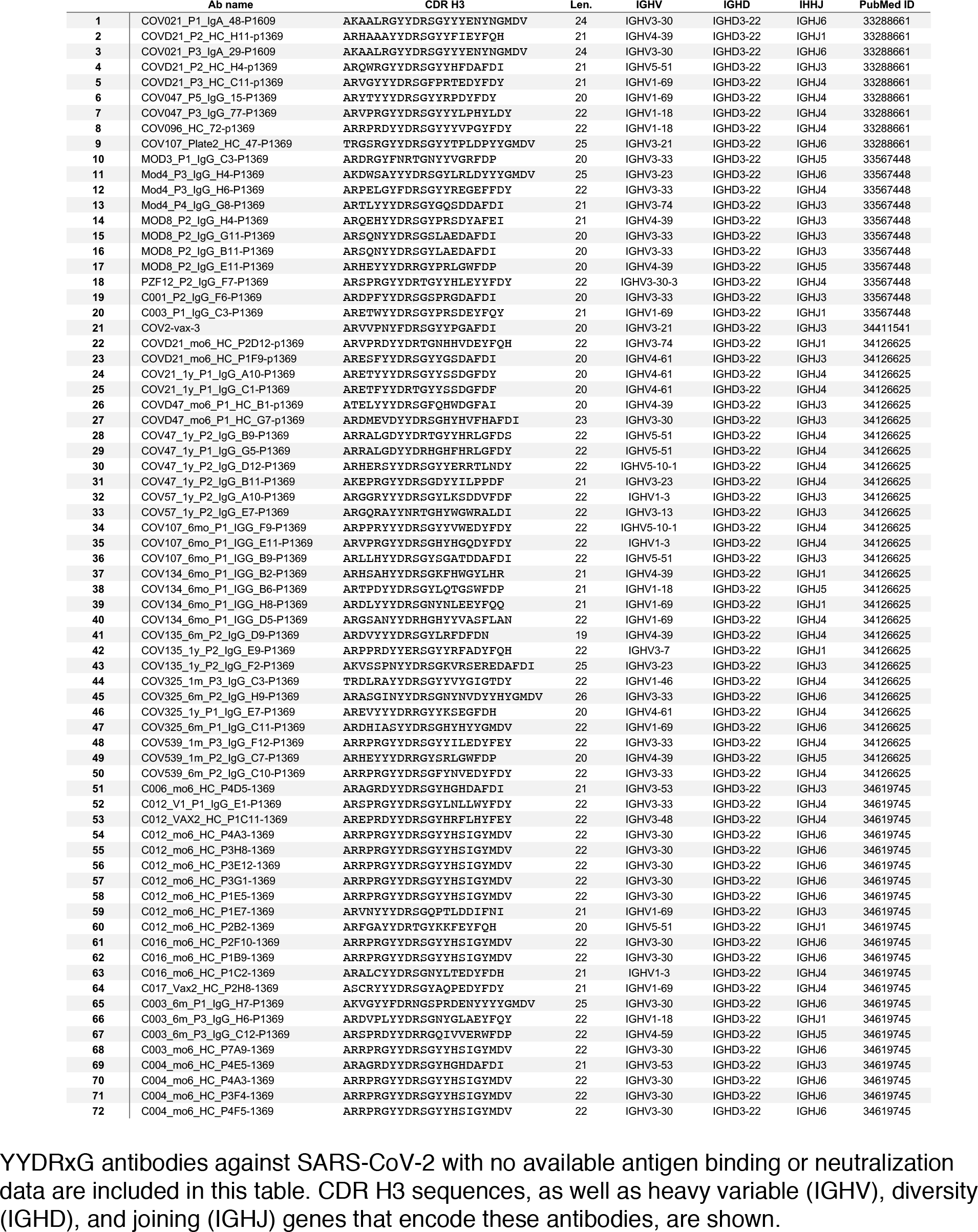
YYDRxG antibodies against SARS-CoV-2 with only sequences available

**Table S4.**
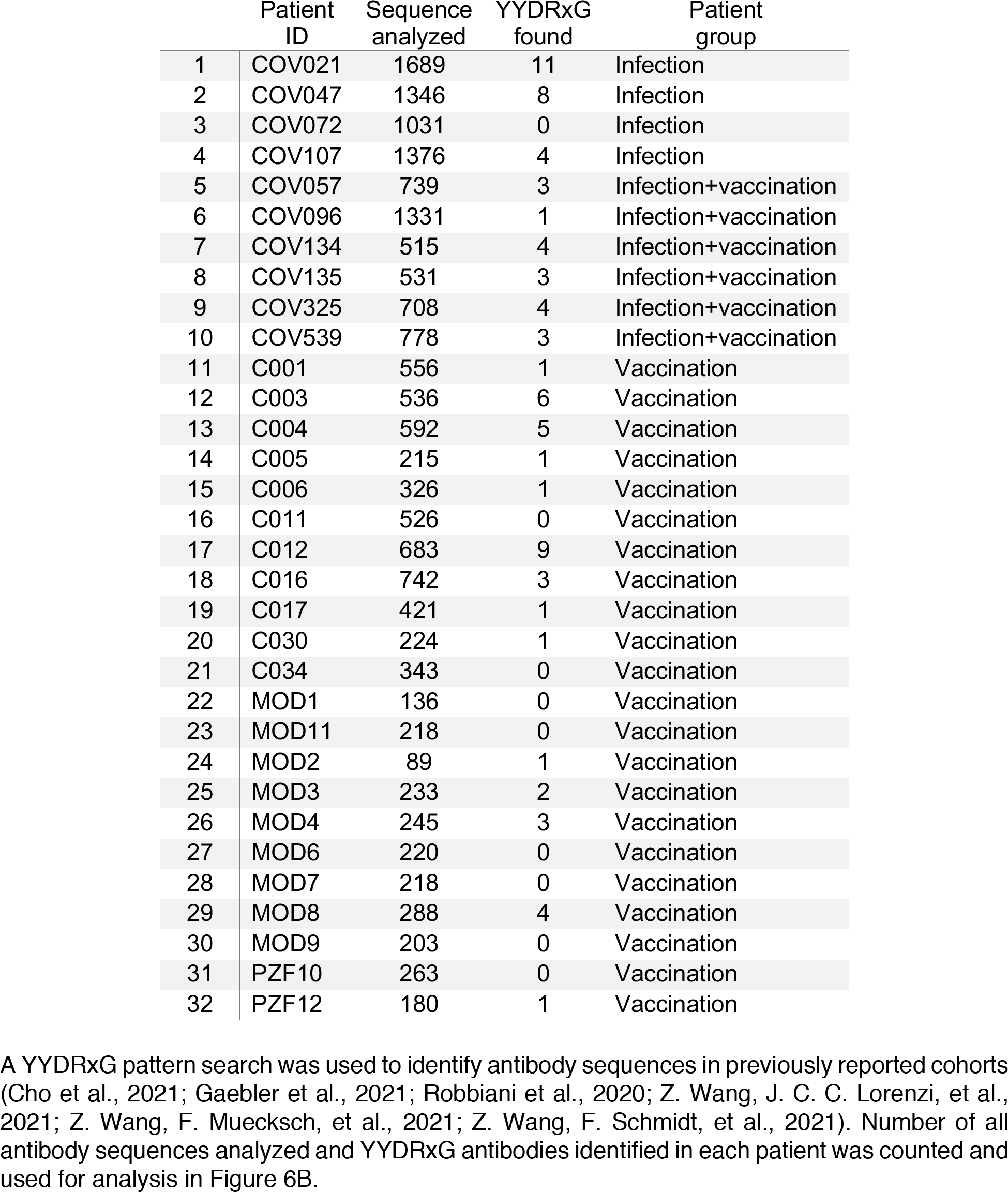
YYDRxG antibodies sequenced from COVID-19 patients and vaccinees

|  | Patient ID | Sequence analyzed | YYDRxG found | Patient group |
| --- | --- | --- | --- | --- |
| 1 | COV021 | 1689 | 11 | Infection |
| 2 | COV047 | 1346 | 8 | Infection |
| 3 | COV072 | 1031 | 0 | Infection |
| 4 | COV107 | 1376 | 4 | Infection |
| 5 | COV057 | 739 | 3 | Infection+vaccination |
| 6 | COV096 | 1331 | 1 | Infection+vaccination |
| 7 | COV134 | 515 | 4 | Infection+vaccination |
| 8 | COV135 | 531 | 3 | Infection+vaccination |
| 9 | COV325 | 708 | 4 | Infection+vaccination |
| 10 | COV539 | 778 | 3 | Infection+vaccination |
| 11 | C001 | 556 | 1 | Vaccination |
| 12 | C003 | 536 | 6 | Vaccination |
| 13 | C004 | 592 | 5 | Vaccination |
| 14 | C005 | 215 | 1 | Vaccination |
| 15 | C006 | 326 | 1 | Vaccination |
| 16 | C011 | 526 | 0 | Vaccination |
| 17 | C012 | 683 | 9 | Vaccination |
| 18 | C016 | 742 | 3 | Vaccination |
| 19 | C017 | 421 | 1 | Vaccination |
| 20 | C030 | 224 | 1 | Vaccination |
| 21 | C034 | 343 | 0 | Vaccination |
| 22 | MOD1 | 136 | 0 | Vaccination |
| 23 | MOD11 | 218 | 0 | Vaccination |
| 24 | MOD2 | 89 | 1 | Vaccination |
| 25 | MOD3 | 233 | 2 | Vaccination |
| 26 | MOD4 | 245 | 3 | Vaccination |
| 27 | MOD6 | 220 | 0 | Vaccination |
| 28 | MOD7 | 218 | 0 | Vaccination |
| 29 | MOD8 | 288 | 4 | Vaccination |
| 30 | MOD9 | 203 | 0 | Vaccination |
| 31 | PZF10 | 263 | 0 | Vaccination |
| 32 | PZF12 | 180 | 1 | Vaccination |

## Notes

### Competing Interest Statement

C.I.K. and L.M.W. are employees of Adimab, LLC and hold shares in Adimab, LLC. L.M.W. is an
315 employee of Adagio Therapeutics Inc. and holds shares in Adagio Therapeutics Inc. L.M.W. is an
316 inventor on a patent describing the ADI anti-SARS-CoV-2 antibodies.

